# Insights from a survey-based analysis of the academic job market

**DOI:** 10.1101/796466

**Authors:** Jason D. Fernandes, Sarvenaz Sarabipour, Christopher T. Smith, Natalie M. Niemi, Nafisa M. Jadavji, Ariangela J. Kozik, Alex S. Holehouse, Vikas Pejaver, Orsolya Symmons, Alexandre W. Bisson Filho, Amanda Haage

**Affiliations:** Department of Biomolecular Engineering, University of California, Santa Cruz, California, United States; Institute for Computational Medicine & Department of Biomedical Engineering, Johns Hopkins University, Baltimore, Maryland, United States; Office of Postdoctoral Affairs, North Carolina State University Graduate School, Raleigh, North Carolina, United States; Morgridge Institute for Research, Madison, Wisconsin, United States; Department of Biochemistry, University of Wisconsin-Madison, Madison, Wisconsin, United States; Department of Biomedical Sciences, Midwestern University, Glendale, Arizona, United States; Division of Pulmonary & Critical Care Medicine, Department of Internal Medicine, University of Michigan, Ann Arbor, Michigan, United States; Department of Biochemistry & Molecular Biophysics, Washington University School of Medicine, St. Louis, Missouri, United States; Department of Biomedical Informatics & Medical Education & The eScience Institute, University of Washington, Seattle, Washington, United States; Department of Bioengineering, University of Pennsylvania, Philadelphia, Pennsylvania, United States; current address: Max Planck Institute for Biology of Ageing, Cologne, Germany; Department of Biology, Rosenstiel Basic Medical Science Research Center, Brandeis University, Waltham, Massachusetts, United States; Department of Biomedical Sciences, University of North Dakota, Grand Forks, North Dakota, United States

**Keywords:** Postdoctoral, Preprints, Mentoring, Funding, Tenure-Track, Faculty Job Search

## Abstract

Applying for a faculty position is a critical phase of many postdoctoral careers, but most postdoctoral researchers in STEM fields enter the academic job market with little knowledge of the process and expectations. A lack of data has made it difficult for applicants to assess their qualifications relative to the general applicant pool and for institutions to develop effective hiring policies. We analyzed responses to a survey of faculty job applicants between May 2018 and May 2019. We establish various background scholarly metrics for a typical faculty applicant and present an analysis of the interplay between those metrics and hiring outcomes. Traditional benchmarks of a positive research track record above a certain threshold of qualifications were unable to completely differentiate applicants with and without offers. Our findings suggest that there is no single clear path to a faculty job offer and that metrics such as career transition awards and publications in high impact factor journals were neither necessary nor sufficient for landing a faculty position. The applicants perceived the process as unnecessarily stressful, time-consuming, and largely lacking in feedback, irrespective of a successful outcome. Our findings emphasize the need to improve the transparency of the faculty job application process. In addition, we hope these and future data will help empower trainees to enter the academic job market with clearer expectations and improved confidence.

## Introduction

The number of doctoral degrees (PhDs) awarded across the globe has increased dramatically in most STEM fields in the past three decades (1,2). The number of faculty positions available has essentially remained stagnant (3), specifically in the United States since 2003 when the National Institutes of Health (NIH) received their last major budget increase (4). Not only are there an insufficient number of faculty positions for the number of PhDs produced (5,6), but trainees typically emerge from academic training feeling underprepared and under-mentored to undertake any other type of job search (7). Hence, there are a large number of applicants per position, many of whom are uncertain about their chances of obtaining a faculty job offer (8,9).

The applicant pool has changed in many ways. PhD graduates are now from a wider range of demographics compared to previous decades, and more diverse than the current hiring committees. Though this is a trend, academic environments are still not representative of the whole population and there are strong calls to push diversity initiatives further (4,10,11). Scientific publishing is also faster-paced with the curriculum vitae (CV) of applicants now looking very different than even 10 years ago. In one study, successfully recruited evolutionary biologists for the title of “junior researchers” needed to publish nearly twice as many articles to be hired in 2013 (22 ± 3.4) than in 2005 (12.5 ± 2.4) (12). The study also found that the length of academic training (defined by them as the time between the first publication and recruitment as a faculty member) has also increased from 3.25 (± 0.6) years in 2005 to 8.0 (± 1.7) years in 2013 (12). This increased length in training time has been reported repeatedly in many STEM fields, and is perceived as detrimental to both the greater scientific community and individuals in these temporary postdoctoral positions (6,13-16).

Despite these changes in training times and hiring trends, the academic job search has largely remained the same, resulting in the perception of the search process as an opaque system with no clear standards or guidelines for navigating the lengthy application procedure. Beyond the requirement of a doctoral degree and possibly postdoctoral training, faculty job advertisements rarely contain information on specific preferred qualifications. Furthermore, the criteria used to evaluate applicants are typically determined by a small departmental or institutional committee and are neither transparent nor made public. The amount of materials required for faculty job applications is also highly variable among hiring institutions. This places a heavy burden not only on the applicants’ time, performance, and well-being, but also on search committees, who spend long hours poring over numerous documents in hopes of identifying the best candidates (17).

Previous studies agree on a need to increase transparency in career outcomes and hiring practices (18-20). The annual pool of faculty job applicants is large and provides a unique opportunity for examining the application process. We performed an anonymous survey, asking for both common components of research and scholarly activity found on an academic CV, as well as information on applicants’ success through the 2018-2019 job cycle. Here we present qualitative and quantitative data on the process as a whole, including information on number of successful off-site and on-site interviews, offers, rejections, and the lack of feedback.

### The application process

Job applicants start by searching for relevant job postings on a variety of platforms (Table S1). The initial electronic application generally consists of a cover letter addressing the search committee, a teaching philosophy statement, CV, and a research plan (Box 2). The length and content of these materials can vary drastically based on the application cycle, region, institution, or particular search committee. In the current system, the overwhelming expectation is that effective application materials require individual tailoring for each specific institution and/or department to which the applicant is applying. This includes department-specific cover letters (21), but may also involve a range of changes to the research, teaching, and diversity statements. The search committee convenes for a few meetings to shortlist the applicants. Applicants are then contacted for interviews somewhere between one to six months after application materials are due. Although many searches usually consist of an off-site (remote) interview, we found that this was not a standard part of every search as the typical applicant received more on-site interviews than off-site. The on-site interview typically lasts 1-2 days and consists of a research seminar, likely a “chalk-talk” (Box 2), and possibly a teaching demonstration. The on-site interview also usually consists of many one-on-one meetings with other faculty members, including a meeting with the hiring department chair, trainees, and the administrative staff. After the interviews, candidates may be contacted and offered a position (Box 2). The time to offer is also variable, but is usually shorter than the time between application and first contact. Importantly, a single search can result in multiple offers (e.g. the department may be able to fund multiple competitive candidates, or the first-choice candidate may decline and the second candidate is given an offer). Searches can also “fail” if the committee does not find a suitable candidate for their program/department or “go dry” if the applicant(s) deemed qualified by the search committee decline their offers. Most applicants are not notified of a receipt of their application, nor are they updated on its status, given a final notice of rejection, or informed that the search may have failed. This uncertainty further complicates an already stressful process that can be mitigated by improving practices for a more streamlined application process.

**Box 1.**
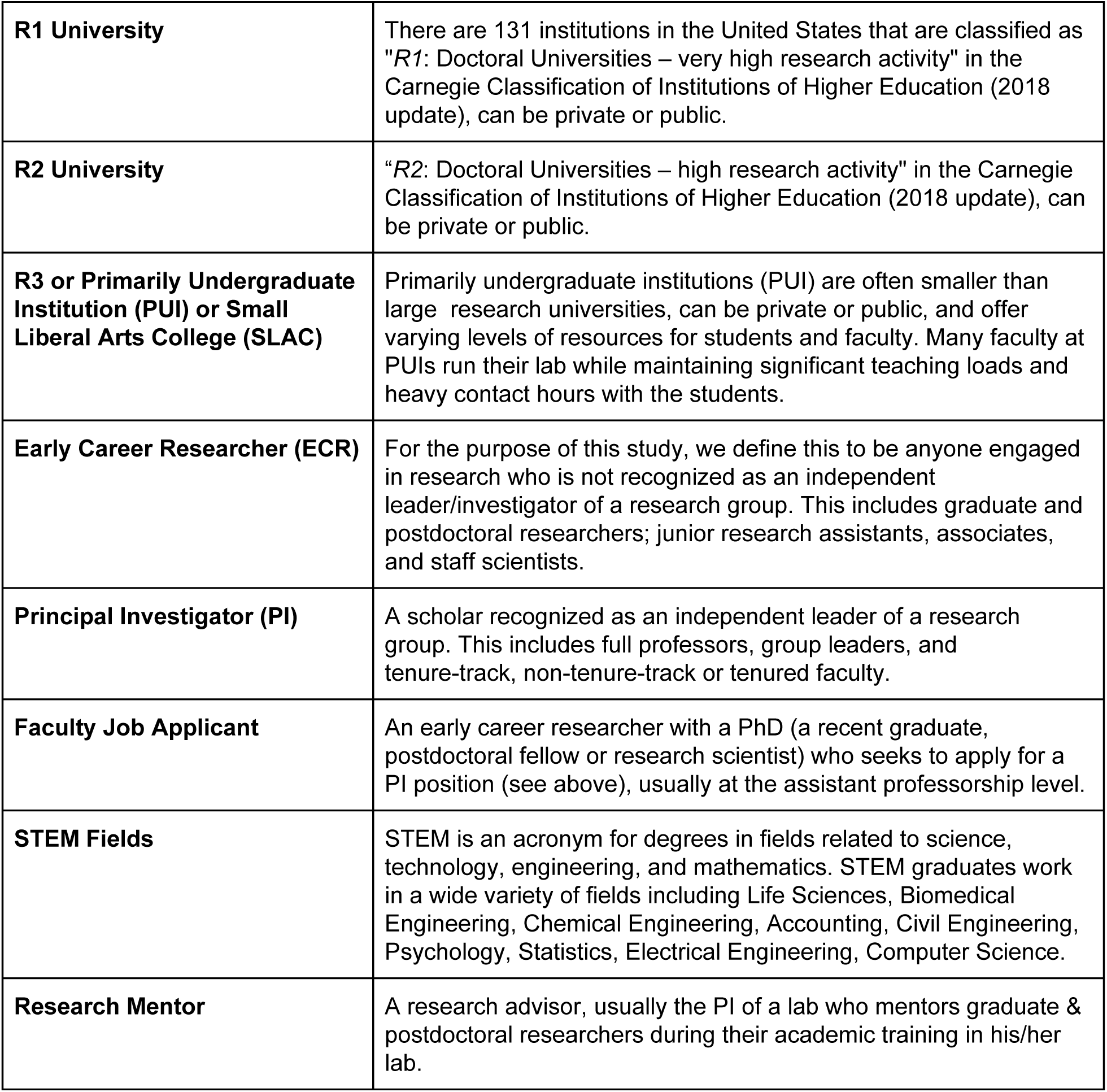
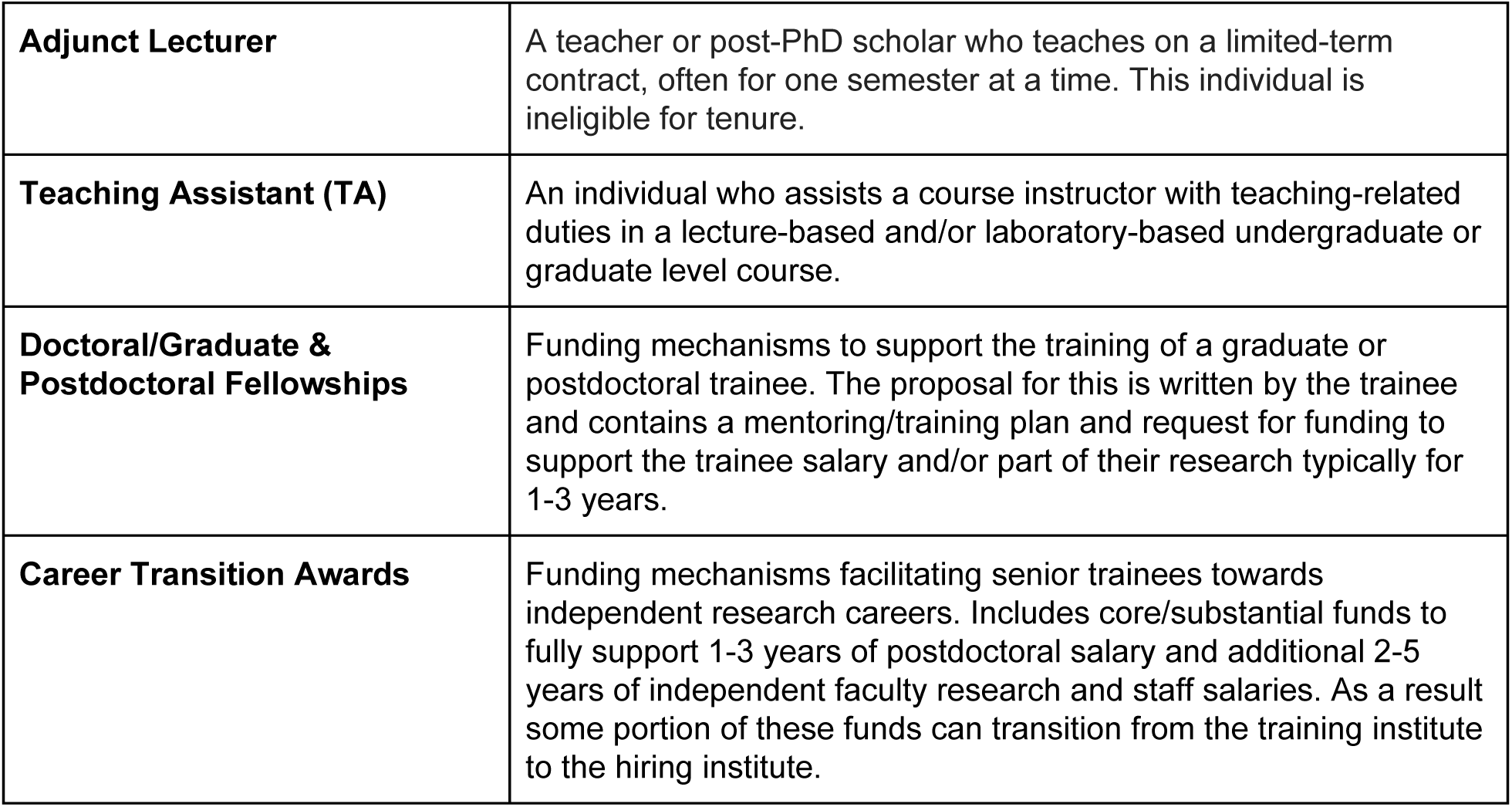
Definition of specific terms used in this study.

**Box 2.**
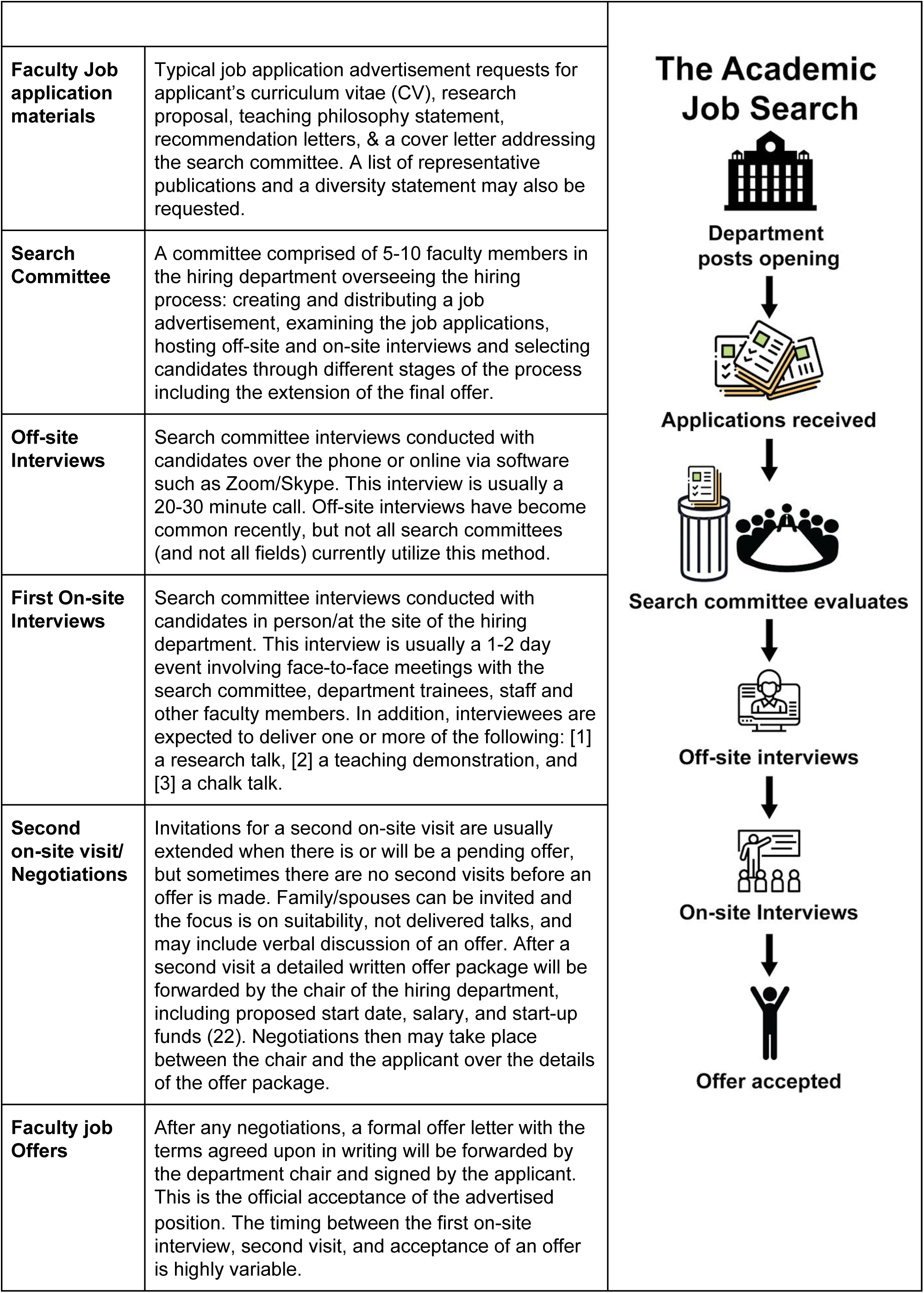
An overview of the Academic Job search process.

| Box 2. An overview of the Academic Job search process |  |  |
| --- | --- | --- |
| **Faculty Job application materials** | Typical job application advertisement requests for applicant's curriculum vitae (CV), research proposal, teaching philosophy statement, recommendation letters, & a cover letter addressing the search committee. A list of representative publications and a diversity statement may also be requested. | The Academic Job Search<br>Department posts opening<br>↓<br>Applications received<br>↓<br>Search committee evaluates<br>↓<br>Off-site interviews<br>↓<br>On-site Interviews<br>↓<br>Offer accepted |
| **Search Committee** | A committee comprised of 5-10 faculty members in the hiring department overseeing the hiring process: creating and distributing a job advertisement, examining the job applications, hosting off-site and on-site interviews and selecting candidates through different stages of the process including the extension of the final offer. |  |
| **Off-site Interviews** | Search committee interviews conducted with candidates over the phone or online via software such as Zoom/Skype. This interview is usually a 20-30 minute call. Off-site interviews have become common recently, but not all search committees (and not all fields) currently utilize this method. |  |
| **First On-site Interviews** | Search committee interviews conducted with candidates in person/at the site of the hiring department. This interview is usually a 1-2 day event involving face-to-face meetings with the search committee, department trainees, staff and other faculty members. In addition, interviewees are expected to deliver one or more of the following: [1] a research talk, [2] a teaching demonstration, and [3] a chalk talk. |  |
| **Second on-site visit/ Negotiations** | Invitations for a second on-site visit are usually extended when there is or will be a pending offer, but sometimes there are no second visits before an offer is made. Family/spouses can be invited and the focus is on suitability, not delivered talks, and may include verbal discussion of an offer. After a second visit a detailed written offer package will be forwarded by the chair of the hiring department, including proposed start date, salary, and start-up funds (22). Negotiations then may take place between the chair and the applicant over the details of the offer package. |  |
| **Faculty job Offers** | After any negotiations, a formal offer letter with the terms agreed upon in writing will be forwarded by the department chair and signed by the applicant. This is the official acceptance of the advertised |  |

|  |  |
| --- | --- |
|  | position. The timing between the first on-site interview, second visit, and acceptance of an offer is highly variable. |

## Results

### Academic Job Applicant Demographics

A total of 322 early career researchers answered survey questions regarding their experience on the academic job market in the 2018-2019 application cycle and data from 317 of these responses were used for analysis (see supplemental materials). Respondents reported a large range in the number of submitted applications from a minimum of one to a maximum of 250 (median: 15). The respondent pool was notably enriched in applicants who received at least one off-site interview (70%), at least one on-site interview (78%) and at least one offer (58%); this may represent a significant bias towards successful applicants in our sample as a recent study shows that only ∼23% (and declining) of PhDs secure a tenure-track position (23).

Respondents represented researchers in a wide variety of fields, with the largest group (85%) from those in the life sciences and related fields, with relatively equal numbers of applications from men and women across the life sciences (Figure 1A, Table S2). Our survey captured data from an international applicant pool, representing 13 countries (Figure 1B, Table S3). However, the majority (72%) of our respondents reported currently working in the United States, which may reflect the larger circulation of our survey on social media platforms and postdoctoral associations there. Most candidates applied to jobs within the United States (82%), Canada (33%), and the United Kingdom (24%) (Table S4). The large majority (96%) of our applicant survey respondents entered the job market as postdoctoral researchers (Figure 1C, Table S5). The applicants spent 1 to 13 years (median: 4 years) in a postdoctoral position. These data are consistent with a recent report suggesting that postdocs in the United States in the field of biomedical sciences spend an average of 3.6 years in their positions (24).

**Figure 1.**
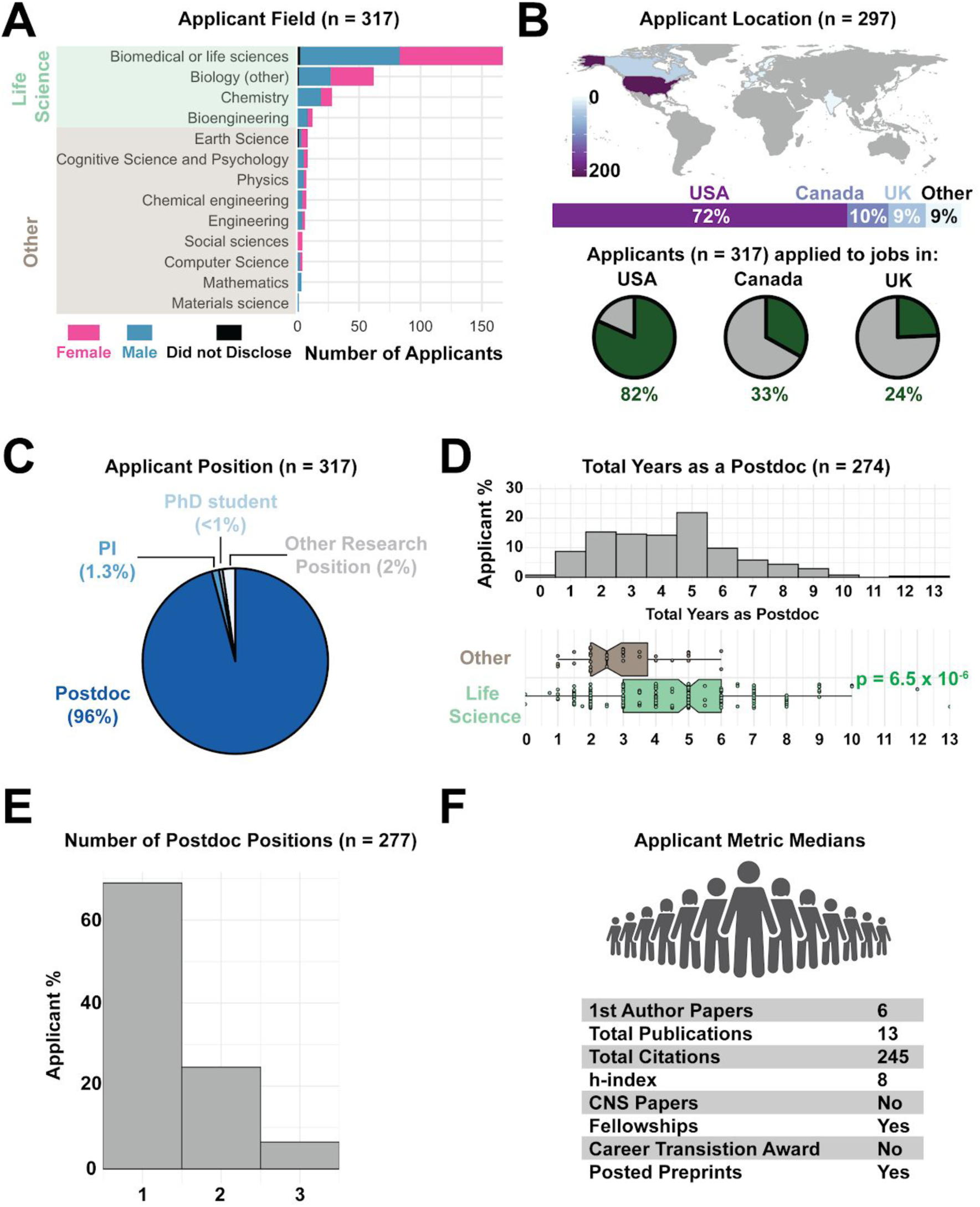
Academic Job Applicant Demographics. A) Distribution of survey respondents by self-identified gender and self-identified scientific field (Table S2). Fields highlighted in green were grouped together as life-science related fields. B) Distribution of the country of residence (research origin) of applicants (top) and the countries in which they applied to for faculty jobs (green slices of pie charts, bottom; for more details see Tables S3 and S34). C) Self-reported positions of applicants when applying for faculty jobs (Table S5). D) The cumulative number of years applicants spent in a postdoctoral fellowship shows that life sciences postdoctoral training takes significantly longer than those in other fields (p-value = 6.5 × 10^−6^) (Table S6, S21). The reported range of years spent as a postdoctoral researcher for our survey applicants was quite large, with 4% of respondents spending 1 year or fewer as a postdoc to 9% of respondents reporting 8 or more years in their postdoctoral positions (maximum of 13 years) (Figure 1D, Table S6). E) Number of postdoctoral positions held by survey applicants (Table S7). F) Median values for metrics of research productivity in the applicant pool.

Notably, postdocs in the life sciences spent significantly more time in postdoc positions (median: 5 years) than those in other fields (median: 2.75 years) before applying for a faculty position (Figure 1D, p = 6.5 × 10^−6^), consistent with previous findings on increased training times in the life/biomedical sciences before junior faculty recruitment (6,12-15). Most survey respondents went on the job market while in their first postdoctoral position (68%) (Figure 1E, Table S7).

Applicants had a large range in their publication records, including number of papers co-authored, h-index, and total citation count. Respondents reported a median of 13 total publications (including co-authorships and lead authorships), with a median of 6 first author papers when entering the job market (Figure 1F, Table S8).

### Applicant scholarly metrics by gender

Gender bias in publishing activity and evaluation is well documented (25-28). Our survey respondents were relatively evenly distributed across self-identified genders, with 51% of applicants identifying as male, 48% as female, while 1% preferred not to disclose this information (with no applicant identifying as non-binary) (Figure 2A, Table S2 first row). Men reported significantly more first author publications than women (medians of 7 and 5, respectively; p = 1.4 × 10^−4^ ; Wilcoxon rank sum), more total publications (medians 16 and 11, p = 3 × 10^−3^) and more overall citations (medians of 343 and 228, p = 1.5 × 10^−2^) (Figure 2B, Table S8, S21). Men in our survey also reported a statistically significant higher h-index than women (medians of 9.0 and 7.0, respectively; p = 5.4 × 10^−3^; Wilcoxon rank sum test) and a higher number of male applicants authored papers in high impact factor journals of *Cell, Nature*, and *Science* (“CNS” journals, 31 women versus 52 men, p = 4.5 × 10^−2^; Wilcoxon rank sum test) (Figure 2C, Table S8, S14, S21). It is of note that applicants self-identified authorship, but the question was clear to include only these three parent journals, excluding other high impact derivatives (see supplemental materials). The gender differences we observe mirror those seen in other reports on differences in citation counts in STEM fields based on the corresponding author gender (29).

**Figure 2.**
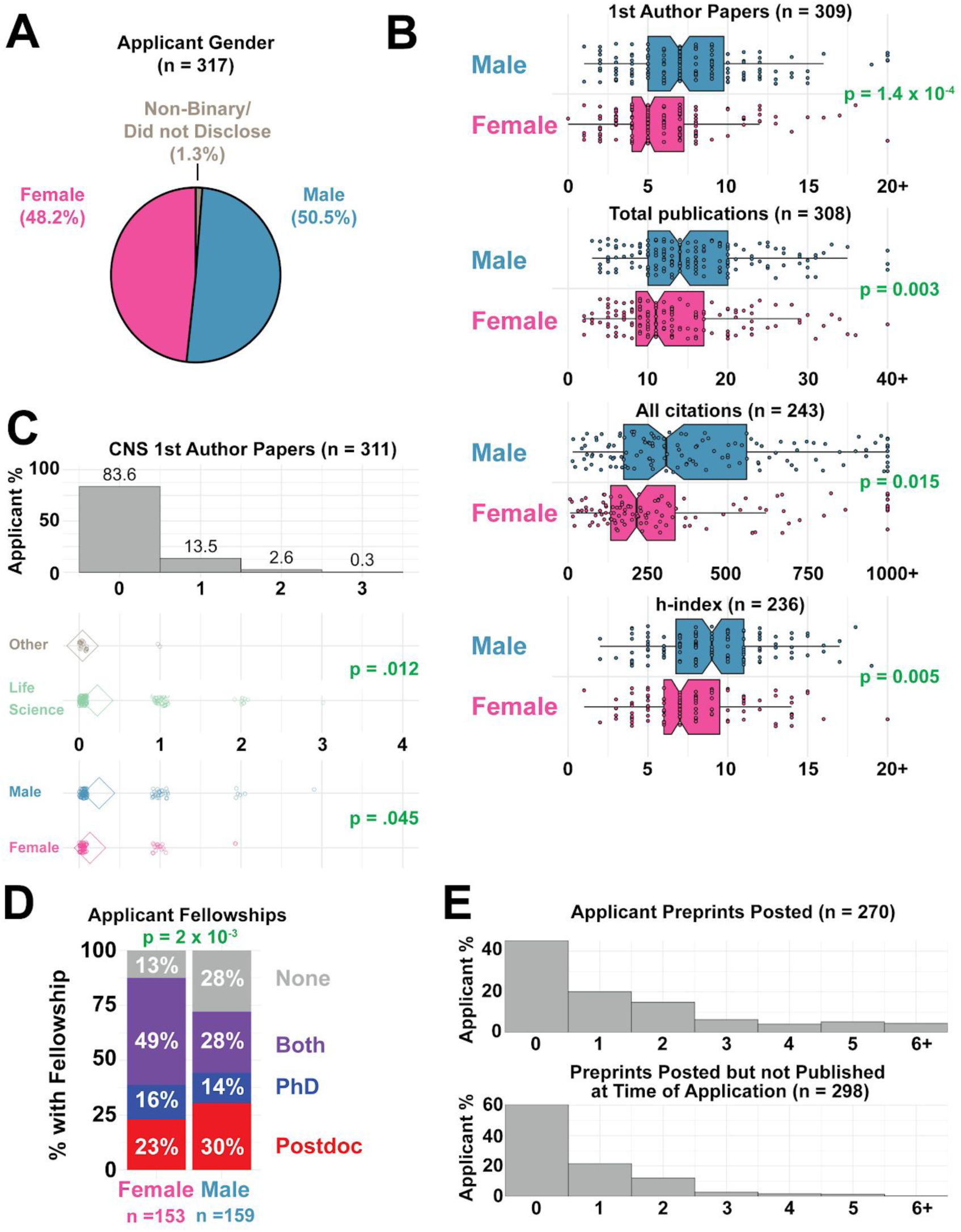
Applicant Scholarly Metrics by gender. A) Distribution of gender (male, female, non-binary, did not disclose) amongst survey respondents (Table S2 first row). B) Publication metrics of survey respondents including number of first author papers (top), total publications (middle top), total citations (middle bottom), and h-index (bottom) for male and female respondents. Men in our survey reported a statistically significant higher h-index than women (medians of 9.0 and 7.0, respectively; p = 5.4 × 10^−3^ ; Wilcoxon rank sum test) (Table S8). C) Number of first author CNS (Cell/Nature/Science) papers are significantly increased for applicants in the life sciences (Table S14) (p = 1.2 × 10^−2^, top) and males (p = 4.5 × 10^−2^). D) Distribution of funding reported within training period (doctoral fellowship only in red, postdoctoral fellowship only in blue, fellowships during PhD and postdoc in purple, and no fellowship in gray) shows females had more fellowship funding than males (p = 2.4 × 10^−3^, Table S9, **χ**^2^ = 12.10). We did not specify the nature of the fellowship in the survey question (i.e. funded by government agencies such as the NIH, private foundations or internal departmental or university sponsored fellowships). E) Number of preprints posted by individual candidates (top) and number of preprints posted which were not yet accepted for journal publication while applying for faculty jobs.

Despite popular discussions on a need for CNS or other high-impact publications (30), the majority (74%) of the applicants who took our survey did not author a CNS paper, and a greater majority (∼84%) did not have a first author publication in a CNS journal (Figure 2C, Table S14). Of the 51 respondents with CNS papers, 49 (96%) were in a life science-related field, indicating the valuation of these journals was highly field-specific (Figure 2C, p = 1.2 × 10^−2^; Wilcoxon rank sum test). While both genders report having obtained fellowships (78%) at some point in their early career, a greater percentage of women received a fellowship of some kind (see Box 1) at some point in their career (p = 2.4 × 10^−3^, **χ**^2^ = 12.10; Chi-squared test, df=2) (Figure 2D). Specifically, 88% of women received a fellowship of some kind (Box 1), compared to 72% of men; women had better success at receiving both doctoral fellowships (42% of women vs 36% of men), and postdoctoral fellowships (72% of women, 58% of men). However, our survey questions did not distinguish between the types (e.g. government funded versus privately funded, full versus partial salary support) or number of fellowships applied to; many of these factors are likely critical in better understanding gender differences in fellowship support. (Figure 2D, Table S9).

### Job application benchmarks and their impact on success

In aggregate, respondents (n = 317) submitted a total of 7,644 job applications (median: 15, average: 24) (Figure 3A, Table S11). Applicants were then invited for a total of 805 (median: 1) off-site (phone, Zoom or Skype) and 832 (median: 2) onsite (campus) interviews, and received 359 offers (median: 1) (Figure 3A, Table S11). In our dataset, 42% of participants received no offers, 33% received one offer, 14% received two offers, 6% received three offers, and 6% received more than three offers. Candidates who received offers typically submitted more applications than those who received no offers, indicating that some candidates may not have submitted enough applications to have a reasonable chance of getting an offer (Figure 3D). A recent Twitter poll with over 700 respondents confirmed that most faculty only received one to three offers during their academic job search (31) (Table S12).

**Figure 3.**
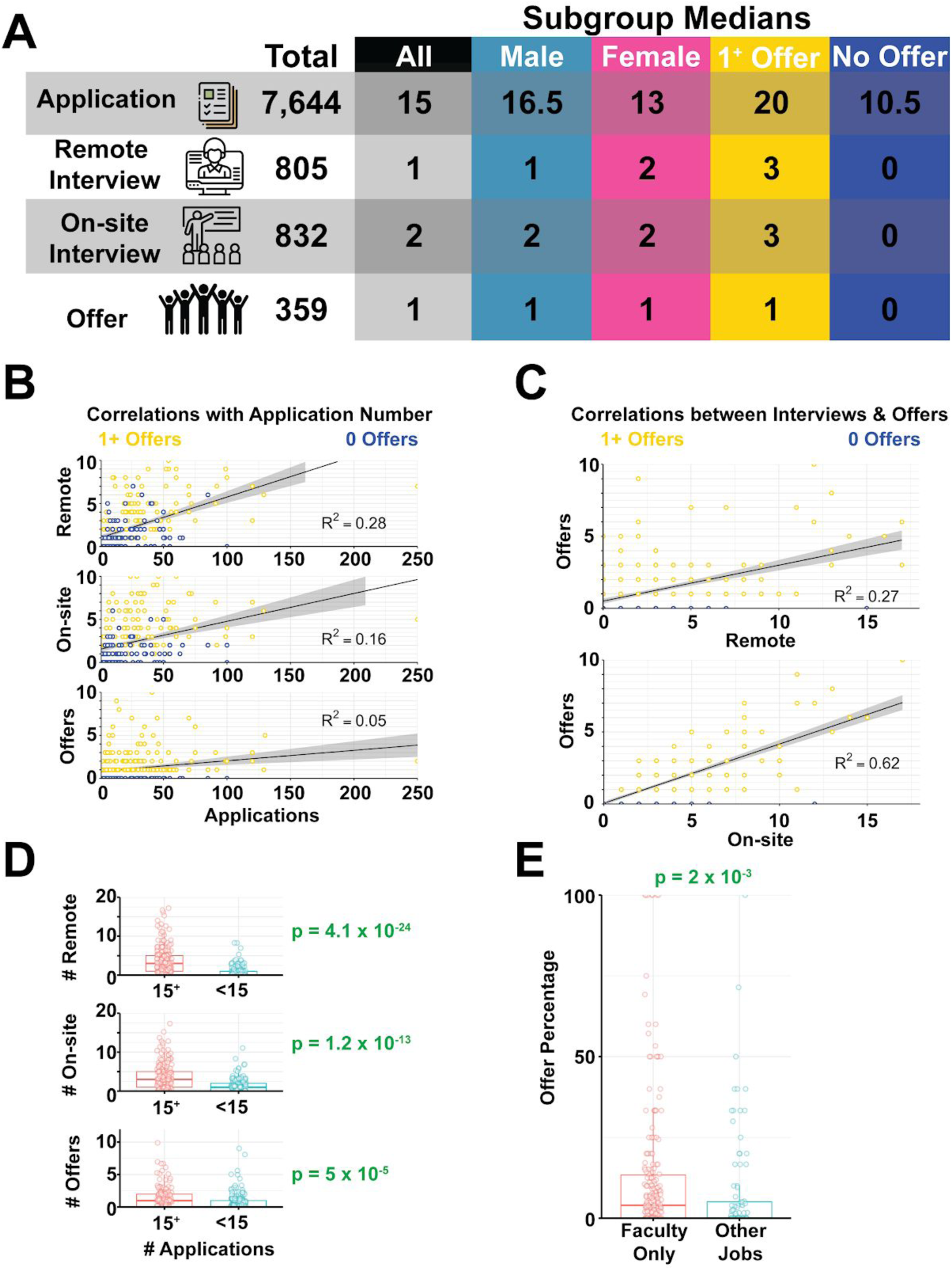
Job application benchmarks and their impact on success. A) Total and median numbers of applications, off-site interviews, on-site interviews and offers recorded in survey responses (Table S8, S11). B) Correlations between the total number of applications submitted and off-site interviews (top), onsite interviews (middle) and offers (bottom). C) Correlations between the number of interviews completed and offers received. See Figure S1 for more details. D) Total number of off-site interviews (top, p < 4.1 × 10^−24^), on-site interviews (middle, p = 1.2 × 10^−13^) and offers (bottom, p = 5 × 10^−5^) for applicants who submitted at least 15 (the median) applications (in red) and less than 15 applications (in blue). E) Fraction of applications that resulted in offers (offer percentages) for survey respondents who did not apply for jobs outside of faculty positions is significantly higher (p = 2 × 10^−3^, Table S21) than for those who also applied for both academic and other types of jobs.

Despite the fact that successful candidates submitted more applications, the number of applications per candidate did not correlate with the number of offers (R^2^ = 4.77 x 10^−2^), while being weakly correlated with the number of off-site interviews (R^2^ = 0.2751) (Figure 3B).

Not surprisingly, the number of on-site interviews strongly correlated with the number of offers received (R^2^ = 0.6239) (Figure 3C). Population medians changed slightly by gender as men submitted slightly more applications, but received slightly fewer off-site interviews. These small differences by gender were not statistically significant (applications p = 7.25 × 10^−2^, off-sites p = 0.1479, on-sites p = 0.5813; Wilcoxon rank-sum test). The median number of offers also did not vary by gender (p = 0.1775; Wilcoxon rank-sum test).

We split our population into two groups by application number, one group either at or below the median (≤ 15 applications, n = 162) and the other group above the median (> 15 applications, n = 155). These groups had a significant difference in success rates: respondents who submitted greater than 15 applications had a significantly higher average number of off-site interviews (p < 4.1 × 10^−24^ ; Wilcoxon rank-sum test), on-site interviews (p = 1.2 × 10^−13^ ; Wilcoxon rank-sum test), and offers (p = 5 × 10^−5^ ; Wilcoxon rank-sum test; Figure 3D). We also asked whether survey respondents applied for non-faculty positions during this cycle (Table S13). Seventy one percent of applicants did not apply for other jobs and they had a small, but significant increase in offer percentage (Figure 3E, p = 1.9 × 10^−3^). Taken together this data seemingly indicates that increasing the number of applications submitted can lead to more interviews, as suggested by others (32), with the typical candidate submitting at least 15 applications to achieve one offer. However, the lower correlation between application number and offers (compared to application number and interviews) suggests that while higher application numbers can generate more interview opportunities, other criteria (e.g. the strength of the interview) are important in turning an interview into an offer.

### Preprints

Preprints, or manuscripts submitted to an open-access server prior to peer-reviewed publication, are becoming increasingly popular among early career researchers (33), particularly in the life sciences, and can boost article citations and mentions (34-38). We investigated whether this increase in preprint submissions had an impact on the chances of receiving an offer. We received 270 applicant responses on this question on preprints; 55% of respondents (148 candidates) had posted at least one preprint, and 20% had posted between 2-6 preprints (average of 1.57 preprints per person) (Figure 2E, Table S8, S15). At the time of faculty job application, 40% of these respondents had an active preprint that was not yet published in a journal, with an average of 0.69 active preprints per person. A number of candidates commented that preprinted research was enormously helpful and served to demonstrate productivity before their paper was published (Tables S26, S27).

### Publication-related metrics

The number of publications authored and the impact factor of those can be predictive of an early career researcher’s chances of obtaining an independent position (39,40). A common perception is that having CNS (*Cell, Nature*, or *Science*) publications are critical to land a life science faculty position (41-43). Our data demonstrates that a CNS paper is not essential to an applicant receiving a faculty job offer as the majority (74%) of our survey respondents did not have authorship of any kind on a CNS paper (Table S14, Figure 4A), while a majority still received at least one offer (58%) (Table S11). Of our respondents, 16% had first authorship on a CNS paper and had significantly higher percentage of offers per application (Wilcoxon rank sum test; p = 1.5 × 10^−4^, median offer percentages: 11% (1^+^ CNS), 2% (No CNS)) and on-site interviews (p = 2.7 × 10^−4^, median onsite 21% (1^+^ CNS), 10% (No CNS)) (Figure 4A). As the number of on-site interviews and offers are highly correlated (Figure 3C), it is unclear if this increased success simply represents a higher chance at landing more onsite interviews. It is important to note that this effect is correlative and these candidates likely had other attributes that made them appealing to the search committee(s).

**Figure 4.**
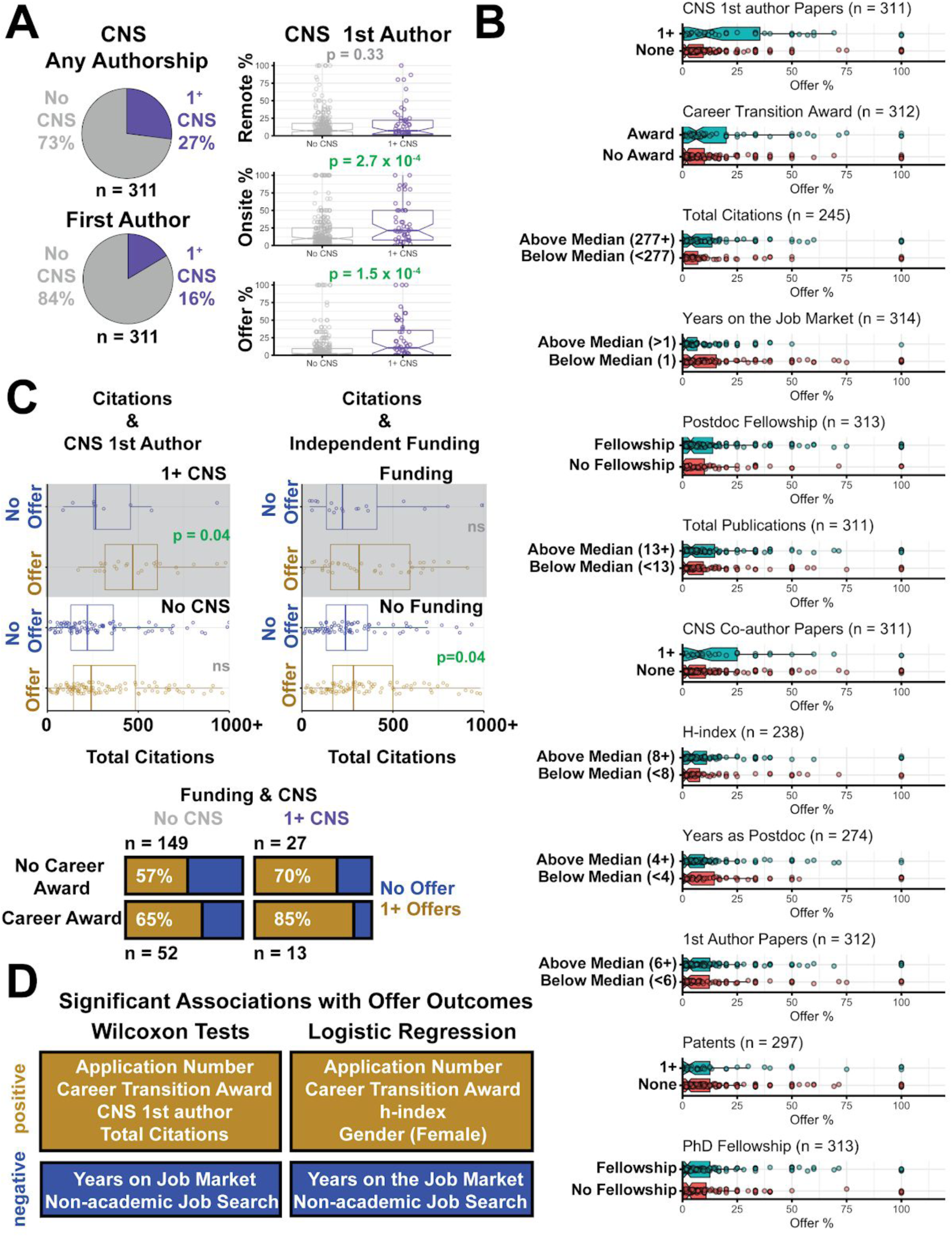
Traditional research track record metrics slightly impact job search success. A) (top left) Fraction of candidates with authorship (purple) of any kind on a CNS paper versus those that did not have CNS authorship (gray). (bottom left) Fraction of candidates with first authorship (purple) on a CNS paper versus those that did not have CNS first authorship (gray). (right) Distributions of off-site interviews (top) (p = 0.33), onsite interviews (middle) (p = 2.7 × 10^−4^) and offers (bottom) (p = 1.5 × 10^−4^) for applicants without a CNS first author paper (“No CNS”, gray), and those with 1 or more CNS papers (“1^+^ CNS”, purple); (Table S14). B) Significant associations were found between offer percentage and the number of first author CNS publications (top, p = 1.7 × 10^−3^), career transition awards (2nd panel, p = 2.5 × 10^−2^), total citations (3rd panel, p = 2.92 × 10^−2^), years on the job market (4th panel, p = 3.45 × 10^−2^). No significant associations between offer percentage were found with having a postdoc fellowship (5th panel), being above the median in the total number of publications (6th panel), having a co-authorship on CNS papers (6th panel), h-index (7th panel), years as a postdoc (8th panel), number of first author papers (9th panel), number of patents (10th panel), and graduate school fellowship status (bottom panel) (Table S8). C) Breakdown of respondents by significant criteria. Citation count (all publications) distributions for groups broken down by CNS 1st authors, career transition awards, and offers (no offer in blue, offer in gold) (top) (p-value = 4 × 10^−2^). Pie-charts: Breakdown of respondents with career transition awards into groups with/without CNS papers and offers (bottom panel) (for applicants with CNS, p = 0.56, **χ**^2^ = 0.34) (for applicants without CNS, p = 0.17, **χ**^2^ = 1.92). D) Summary of results testing criteria with offer outcomes either through Wilcoxon analyses or logistic regression.

We examined several other publication metrics and found no correlation with number of offers. Specifically, the total number of publications (R^2^ = 8 × 10^−2^), the number of first author (R^2^= 2 × 10^−2^), the number of corresponding author publications (R^2^ = 9 × 10^−4^), and h-index (R^2^ = 4 × 10^−3^) did not significantly correlate with offer percentage (Figure S1). When we separated candidates who were above and below the medians for each of these metrics and compared the distribution of offer percentages, only the total number of citations significantly associated with a higher offer percentage (p = 2.9 × 10^−2^) (Figure 4B). Although offer percentage was generally higher for applicants above the median for the other metrics, none of these differences were statistically significant (first author publications p = 1.0, total publications p = 0.190, h-index p = 0.57; Wilcoxon rank-sum test with Holm correction) (Figure 4B, Table S21).

### Doctoral and postdoctoral fellowships

Respondents were highly successful in obtaining fellowship funding during their training (80% received a fellowship of any kind, Figure 2D, Tables S9). Applicants with a postdoctoral fellowship had a greater offer percentage than those without, although the effect was not significant after correcting for multiple comparisons (p = 0.169; Wilcoxon rank sum test, Holm correction, Figure 4B). Doctoral fellowships did not appear to influence an applicant’s offer percentage (p = 1.0; Wilcoxon rank sum test, Holm correction, Figure 4B).

### Career transition awards

Receiving funding as an early career researcher is part of a favorable research track record (44). A recent study of publicly available data indicates that the proportion of faculty receiving their first large research program grant (an R01 through the NIH) with a history of funding as a trainee (F and K awards through NIH) is significantly increasing, driven mostly by K awards. Pickett *et al* state that “While not a prerequisite, a clear shift is underway that favors biomedical faculty candidates with at least one prior training award” (45). Our survey differentiated the types of funding a trainee can receive into predoctoral and postdoctoral fellowships (discussed above), and career transition awards, for which the trainee is listed as the PI and funds can often transition with the trainee to a hiring institute (e.g. the Burroughs Wellcome Fund Career Awards at the Scientific Interface or the NIH K99/R00 Pathway to Independence award). Career transition awards were less frequent in our survey responses, with 25% of respondents receiving awards on which they were PI/co-PI (Tables S10). Respondents with transition funding received a higher percentage of offers (p = 2.5 × 10^−2^; Wilcoxon rank sum test, Holm correction, Figure 4B).

### Patents

Patents are considered positive metrics of research track record, although their importance and frequency can vary between fields. Only 19% of applicants reported having one or more patents on file from their work when entering the job market (Table S16). The number of patents held by the applicant did not correlate with the number of offers received (R^2^ = 3 × 10^−3^) (Figure S1) and the percentage of offers did not change between those with or without a patent (p = 1.0; Wilcoxon rank-sum test, Holm correction) (Figure 4B).

### Years on the Job Market

We also asked respondents on the number of application cycles in which they had participated. Roughly half (55%) of our respondents were applying for the first time, and these candidates fared significantly better in terms of offer percentages than those who were applying again (p = 3.5 × 10^−2^ ; Wilcoxon rank-sum test, Holm correction) (Figure 4B). Analyses such as the work presented here may help applicants refine and present their materials and track record in a manner that might improve success and decrease repeated failed cycles for applicants.

### Interplay between metrics

We next examined the relationship between each of the traditional criteria that were significantly associated with an increase in offer percentage (CNS 1st authorship, total citations, and career transition awards). Overall we had 241 applicants that fully responded to all of our questions about these metrics. Pairwise testing of each of these criteria found no statistically significant relationships between variables (p = 0.446, career transition awards vs CNS; p = 0.264 total citations vs CNS; p = 0.289 career transition awards versus total citations; Chi-squared tests). Regardless, we plotted subgroups based on offer status and each of these criteria to see if there was evidence for any general trends in our dataset (Figure 4C). Notably, respondents with a CNS first authorship who received at least one offer had a greater number of total citations than those with a CNS 1st authorship, but no offers (Figure 4C). Applicants with CNS first authorship or career transition awards (Figure 4C) had higher percentages of securing at least one offer, and those with both had an even greater percentage although the differences between these groups was not statistically significant.

This analysis suggests that the combination of different criteria holistically influence the ability to obtain an offer. Therefore, we performed logistic regression to examine the relationship between multiple variables/metrics on the successful application outcome of receiving an offer (Table S38). When applicants with missing values were excluded, only the number of applications was found to be significantly associated with offer status (*β* = 0.4874, p = 4.4 × 10^−2^) among the 105 applicants that remained. When missing values were imputed, significant positive coefficients were observed for h-index (*β* = 0.6709, p = 1.9 × 10^−2^), application number (*β* = 0.6411, p = 1.23 × 10^−4^), career transition awards (*β* = 0.3913, p = 5.86 × 10^−3^) and identifying as female (*β* = 0.3039, p = 3.3 × 10^−2^). Moreover, the search for non-academic jobs (*β* = −0.3477, p = 9.76 × 10^−3^) and the number of years on the job market (*β* = −0.2887, p = 3.7 × 10^−2^) were significantly negatively associated with offer status (Figure 4D, Table S38). We note that the model with imputed data was more accurate than that with missing values excluded at distinguishing between applicants with and without offers in 10-fold cross-validation experiments. However this accuracy was found to only be 65%, which is insufficient to construct a usable classifier of offer status.

Finally, we extended this analysis to visualize the interplay between all variables in Figure 4B by learning a decision tree automatically from the collected data (Figure S3). The algorithm tries to partition the applicants into groups such that each group is entirely composed of individuals with at least one offer or without. A variety of different classifier groups were identified, but no group contained more than ∼19% (61 out of 317) of the dataset. In fact, the accuracy of the overall decision tree in distinguishing between candidates with offers and those without was only ∼59% (see supplemental material). Taken together, these results suggest that there are multiple paths to an offer and that the variables we collected do not sufficiently capture this variability among applicants.

### Most applicants fulfill the teaching requirements for any university type

Discussion surrounding the academic job market is often centered on applicants’ publications and/or funding record, while teaching experience generally receives much less attention. Accordingly, a candidate’s expected teaching credentials and experience varies and largely depends on the type of institution at which the candidates are seeking a position. We limited survey response options to either R1 focused, PUI focused, or submitting applications to both types of institutions (Box 1). A majority of our respondents applied to jobs at R1 institutions (Figure 5A, Table S17), which may be the reason that most discussions focus on research-centric qualifications. Despite this, almost every application to an R1 institution requires a teaching philosophy statement. The majority of our survey respondents (99%) have some type of teaching experience (Figure 5B, Table S18), with roughly half of applicants’ teaching experience limited to serving as a Teaching Assistant (TA) only (Box 1), while the other half reported experience beyond a TA position, such as serving as an instructor of record (Figure 5B, Table S18, S19). The degree of teaching experience did not change based on the target institution of the applicant (p = 0.56, **χ**^2^= 0.41; Chi-squared test, Figure 5C), nor did the percentage of offers received significantly differ between groups based on teaching experience (p = 0.16; Wilcoxon rank-sum test, Figure 5D).

**Figure 5.**
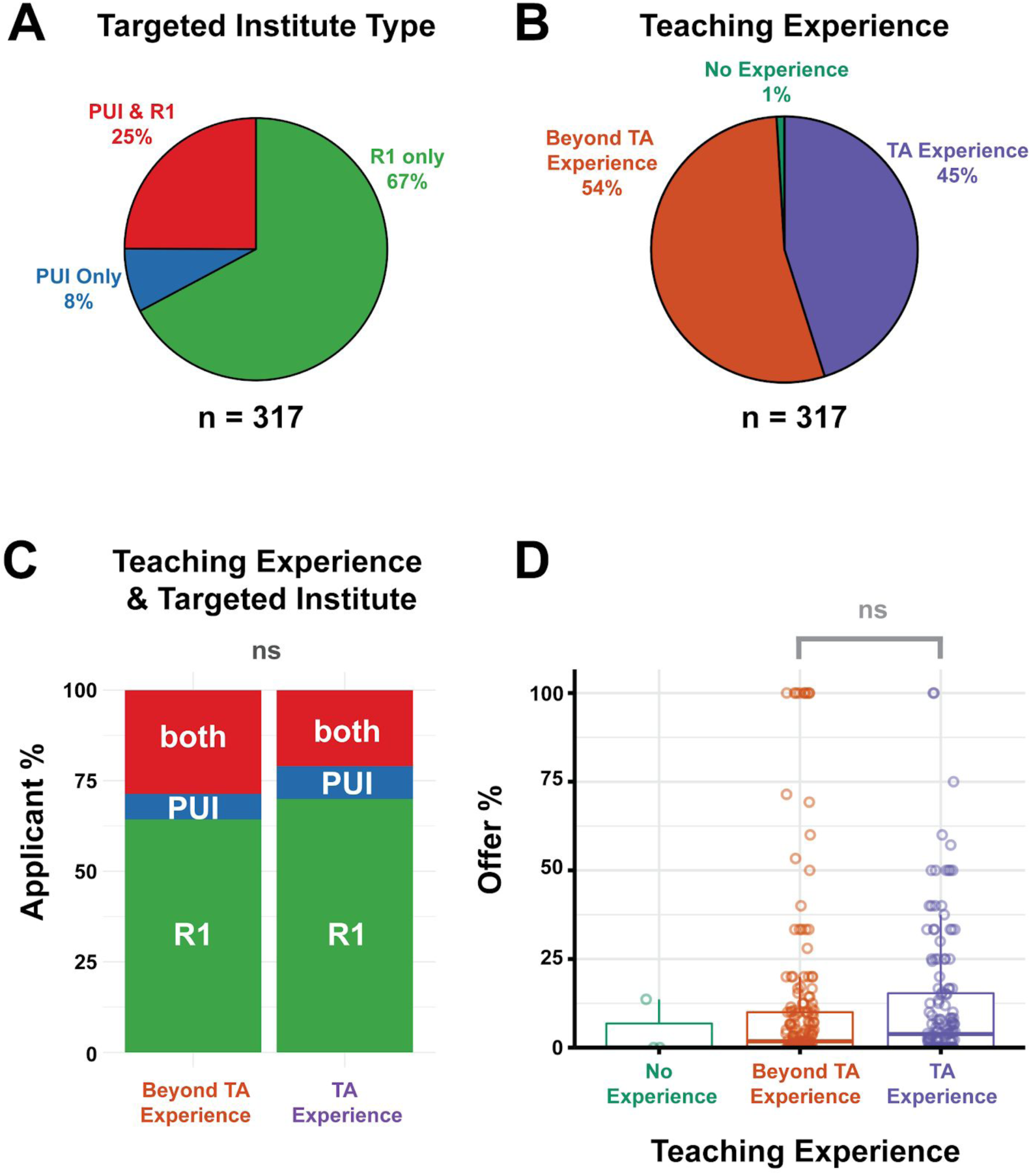
Summary of applicant teaching experience and impact on job search success. A) Distribution of institution types targeted by survey applicants for faculty positions (PUI only (in blue), R1 institutions only (in green) or both (in red), Table S17). B) Distribution of teaching experience reported by applicants as having TA only experience (in purple), beyond TA experience (e.g. teaching certificate, undergraduate and/or graduate course instructorship, guest lectureship and college adjunct teaching, (in orange), or no teaching credentials (in green) (Table S18). C) Distribution of teaching experience (TA experience, right, vs. Beyond TA experience, left) for applicants who applied to R1 institutions only (in green), PU institutions only (blue), or both R1 and PUIs (in red), Table S18, p = 0.56 (ns), **χ**^2^ = 0.41; Chi-squared test). D) Association between offer percentage and teaching experience is not significant (p = 0.16; Wilcoxon rank-sum) (Tables S18-S19, S21).

### Research versus Teaching-intensive institutions

To our knowledge, there is a lack of systematic evidence describing the process or expected qualifications of a PUI-focused (Box 1) job search (46). A subgroup of 25 “PUI only” applicants responded to our survey, and despite this small number, we aimed to describe this important sub-group here. Many characteristics of the whole survey population were reflected in this sub-group, including gender identity, number of publications, funding, and teaching experience (Figure 6A-D). The median number of remote interviews, onsite interviews, and offers was also similar to R1-focused applicants with PUI-focused applicants submitting fewer applications (Figure 6E). Interestingly, when asked to describe their “beyond TA” teaching experience, this sub-group was specifically enriched in “adjunct”, “visiting professor”, “instructor of record”, “community college”, or “contract-based” teaching experiences compared to the ‘R1 only’ or ‘both’ applicant groups (p = 5 × 10^−4^, **χ**^2^ = 27.5; Chi-squared test, Figure 6F, Table S19). Having “adjunct” experience as described in Figure 6F did not significantly increase the median number of offers received for PUI focused applicants (p = 0.55; Wilcoxon rank-sum test, Figure 6G). There was no difference in the median number of offers received based on adjunct experience for applicants targeted at R1s or both types of institutions (p = 0.99; Wilcoxon rank-sum test, Figure 6G).

**Figure 6.**
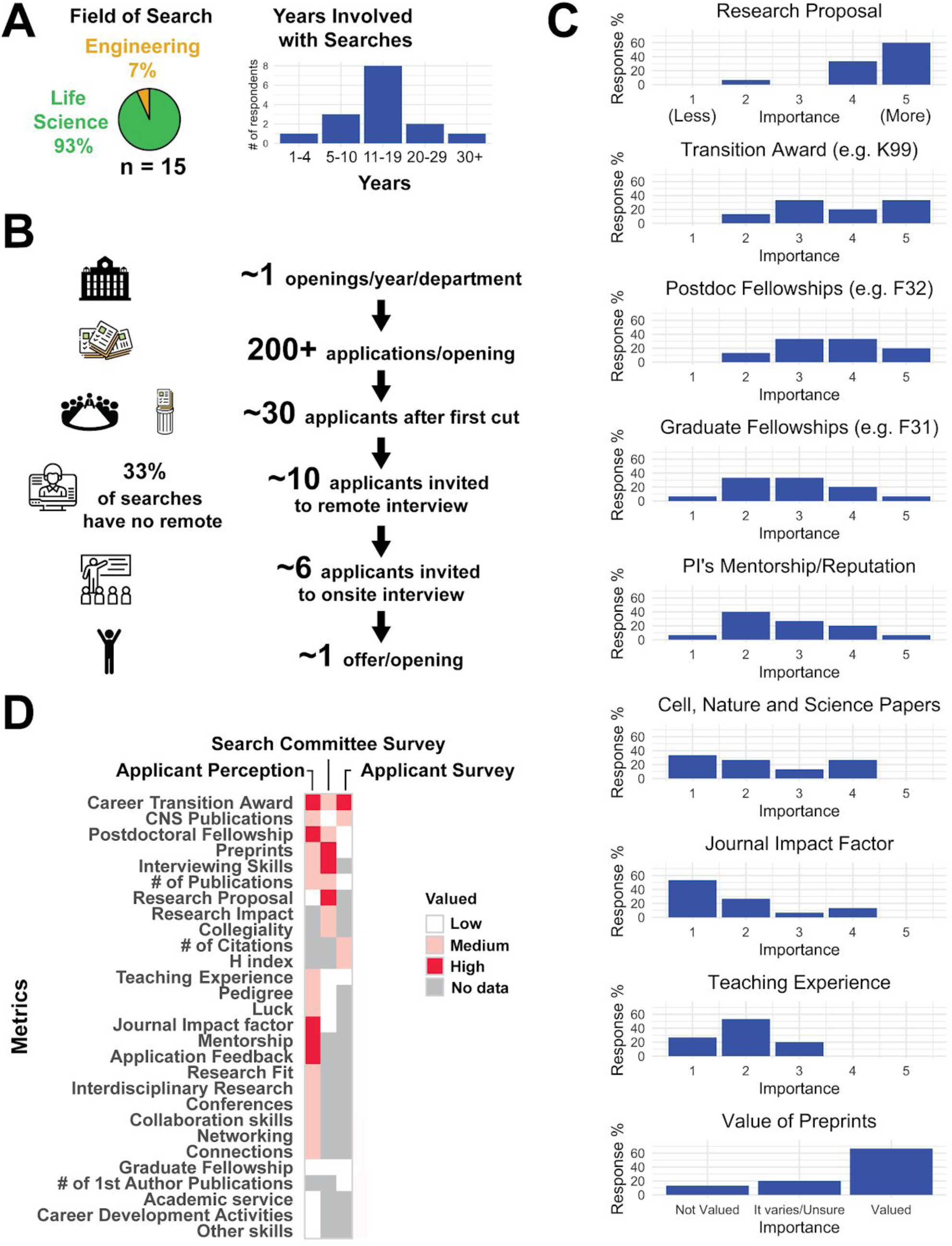
PUI focused applicants differ only in teaching experience from the rest of the application pool. A) The gender distribution of the applicant who focused on applying to PUIs (Table S17). B) The gender distribution and number of first-author publications of the applicant who focused on applying to PUIs (p = 0.88; Wilcoxon rank-sum test). C) Summary of the fellowship history by gender for PUI focused applicants (Table S9). D) Distribution of teaching experience of PUI focused applicants (Table S18). E) The median number of applications, off-site interviews, on-site interviews and offers for PUI focused applicants. F) Percentage of survey respondents who identified having “adjunct teaching” experience (Box 2) based on target institution (p = 5 × 10^−4^ ; **χ**^2^ = 27.5, Chi-squared test). G) The number of offers received segregated by “adjunct” experience in either PUI focused applicants (p = 0.55; Wilcoxon rank-sum test) or R1/both R1 & PUI focused applicants (p = 0.98; Wilcoxon rank-sum test).

### Applicants perceive the process to be time-consuming and opaque, with minimal to no feedback

We asked the applicants to comment on whether any aspect of their training or career was particularly helpful or harmful to their faculty applications (Table S22). We used word clouds and follow-up Twitter polls (Tables S12, S24-25) to analyze recurrent themes in these open-ended questions. The applicants identified “funding” as most helpful for their applications, and “no-funding” as subsequently harmful; this perception agrees with the data presented above (Figures 7A,4C,1S). Additionally, our applicants’ perceptions were also in line with the rest of the data, in that they were unable to largely agree on other measurable aspects of their career that were perceived as helpful. Qualitative aspects that were perceived as particularly helpful included networking and attending/presenting at conferences. Interestingly “interdisciplinary-research”, which is often highlighted as a strength and encouraged by institutions and funders, was perceived by candidates as a challenge to overcome. Indeed, interdisciplinary candidates may pose an evaluation challenge for committees given the differences in valuation of research metrics across fields, the extended training time required to master techniques and concepts in multiple fields, as well as valuation of interdisciplinary teams of specialists over interdisciplinary individuals (47).

**Figure 7.**
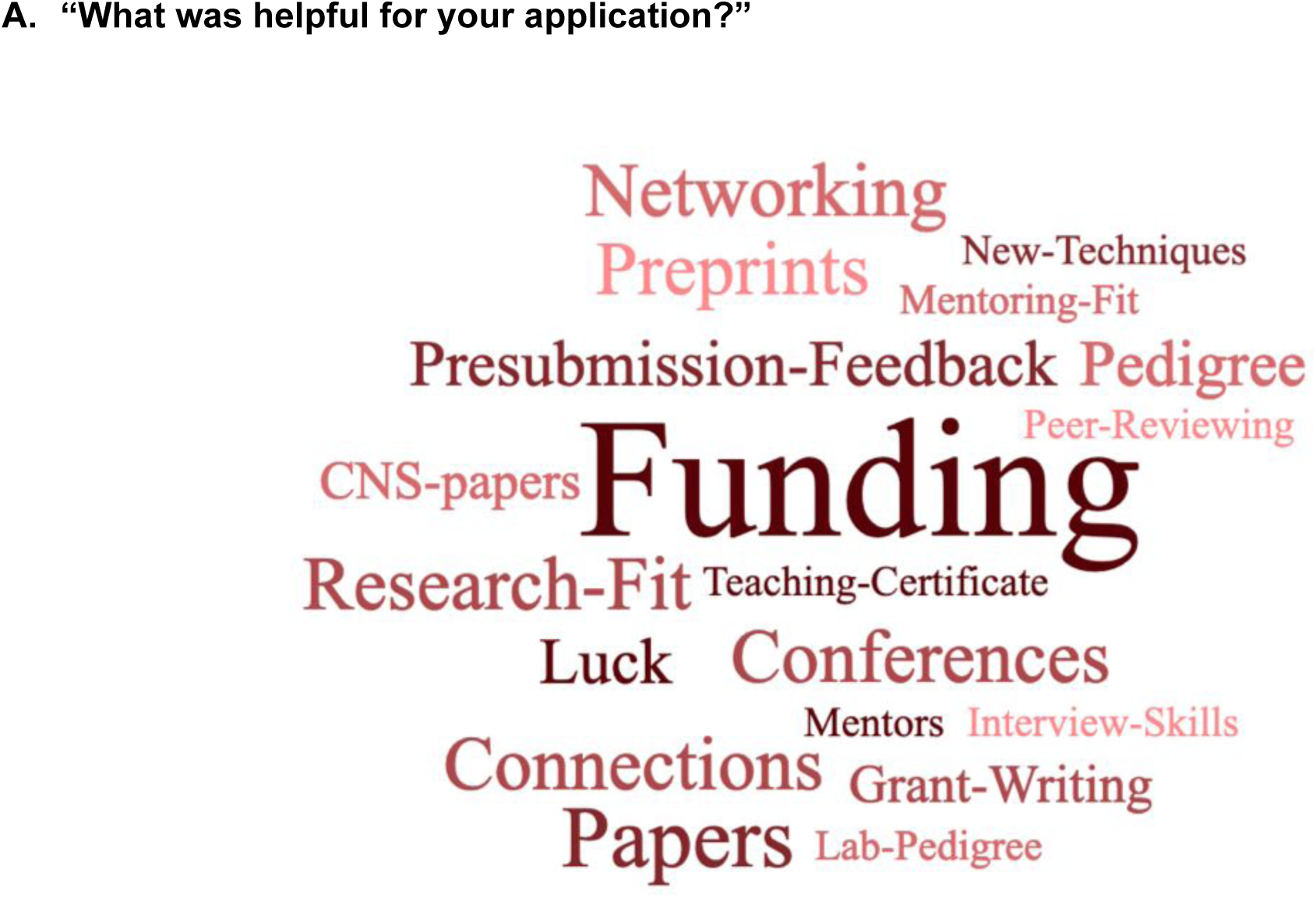
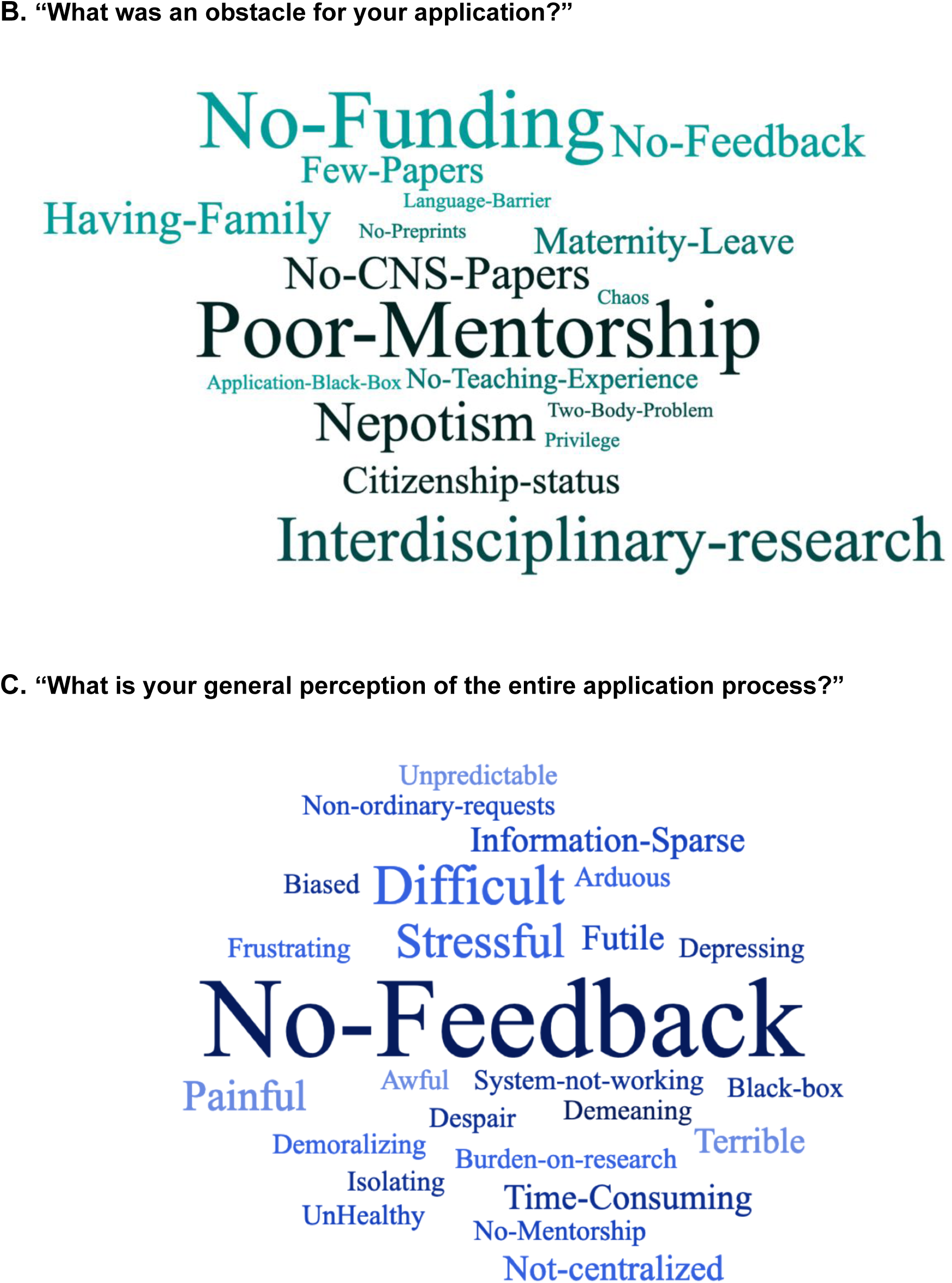
Applicants’ perception of their faculty job application process. Three word clouds summarizing qualitative responses from the job applicant survey respondents to the following questions: A) “What was helpful for your application?” (top) (Table S22), B) “What was an obstacle for your application?” (middle) (Table S26), and C) “What is your general perception of the entire application process?” (bottom) (Table S27). The size of the word (or short phrase) reflects its frequency in responses (bigger word corresponds to more frequency). Survey respondents were able to provide longer answers to these questions, as shown in Tables S22, S26-S27. “CNS-papers” is an abbreviation for Cell/Nature/Science publications, “Pedigree” refers to the applicant’s postdoc lab pedigree or postdoc university pedigree, “Grant-Writing” refers to the applicant’s grant writing experience with their PhD or postdoctoral mentor, “Peer-reviewing” refers to the experience of performing peer-reviewing for journals, “Interdisciplinary-research” refers to comments stating that Interdisciplinary research was underappreciated, “two-body-problem” refers to the challenges that life-partners face when seeking employment in the same vicinity, “No-Feedback” refers to lack of any feedback from the search committees on the status, quality or outcome of applications.

Notably, many applicants found the amount of time spent on applications and the subsequent lack of feedback from searches frustrating (Figure 7B-C, Tables S11,S22,S25). Most applicants never received any communication regarding their various submissions. For instance, an applicant who applied for 250 positions only received 30 rejections. Overall, our pool of applicant survey respondents submitted 7,644 applications (Figure 3A) and did not hear anything back in 4,365 cases (57% of applications), receiving 2,920 formal rejection messages (38% of applications) (Table S11). Application rejection messages (if received at all) most often do not include any sort of feedback. Additionally, a considerable amount of time is spent on writing each application; as previously noted, extensive tailoring is expected for competitive materials. A separate Twitter poll indicated that applicants in general, not specifically for this cycle, typically spend more than 3 hours tailoring each application (48) (Table S24). Our pooled applicants at minimum then spent a combined 22,932 hours (7,644 applications x 3 hours preparation each), or 2.62 years, on these applications. Individually, this number amounts to 72 hours for each applicant on average, but does not take into account how long the initial creation of “base” application materials takes, which is often a much longer process. In another follow-up Twitter poll, a majority of respondents felt that time spent on preparing faculty job applications impeded their ability to push other aspects of their career forward (Table S25) (49). Combining these insights, it is therefore unsurprising that almost all applicants, including applicants that received at least one offer (Table S28), found the process “time-consuming”, a “burden-on-research”, and “stressful” (Figure 7B-C, Table S27, Table S28).

Forty four percent of our respondents had applied for PI jobs for more than one cycle (Table S30). Though applicants who applied for more than one cycle had significantly lower offer percentages (p = 3.45 × 10^−2^; Wilcoxon rank-sum test; Figure 4B), many reported perceived benefits from significant feedback from their current PI through their previous application cycles. Though mentorship was not as often reported as specifically helpful (Table S26), the lack of mentorship was a commonly cited harmful obstacle (Figure 7B, Table S27). Lastly, multiple candidates felt that issues pertaining to family, a two-body problem (need for spousal/significant other hire), or parental leave significantly harmed their success.

### Search Committees Value the Future

In an effort to better understand the academic search process, we performed an exploratory survey of academic search committee members. Fifteen faculty members responded, with 67% having been involved in search committees for over ten years (Figure 8; Table S31). In keeping with the academic and geographical contours of our applicant survey respondents, we focused on faculty members at R1 academic centers working in life sciences (93% of those polled) and engineering (7%) within the United States (Table S32).

**Figure 8.**
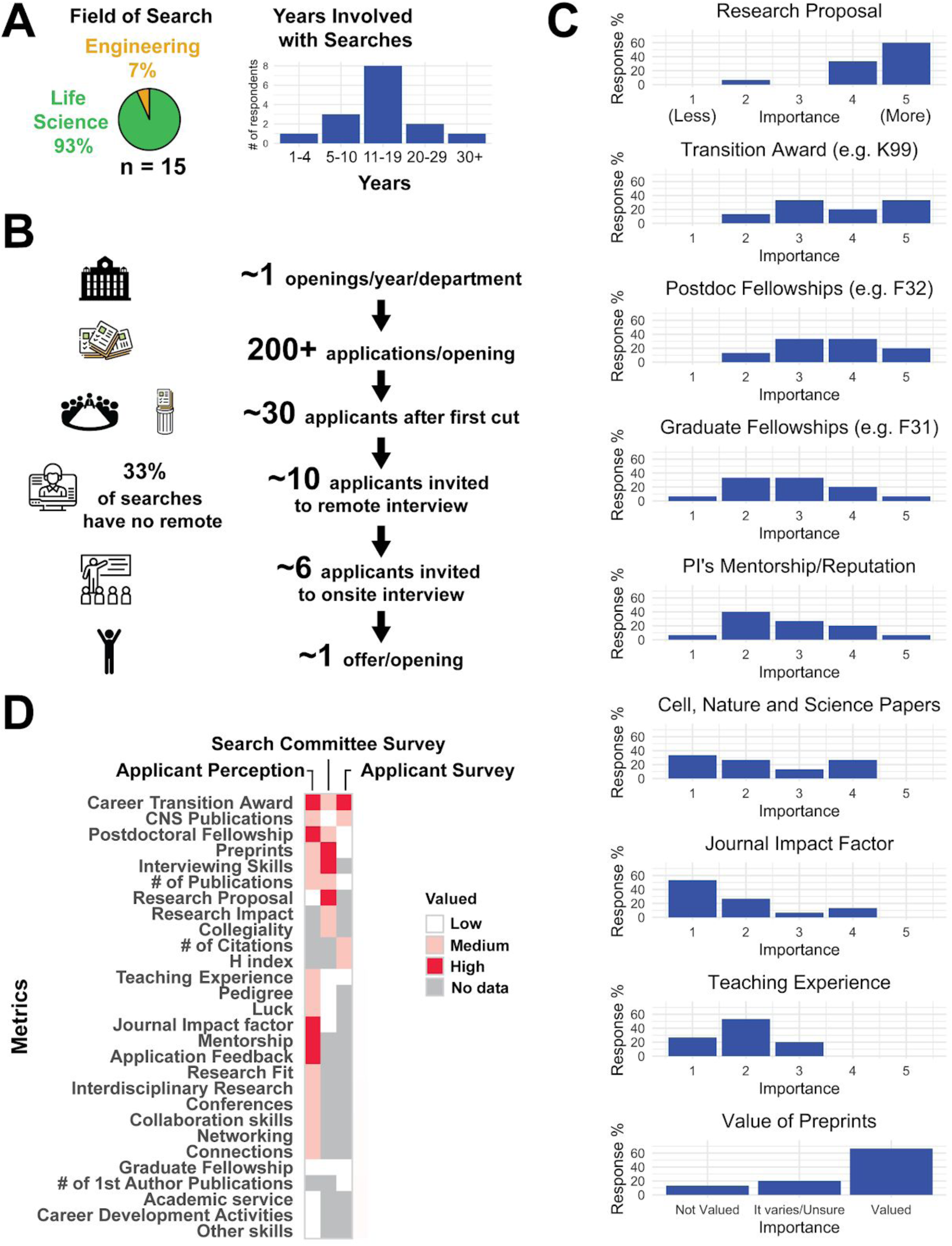
Summary of metrics valued by search committees. Search committee members were asked on how specific factors were weighted in the decision on which applicant to extend an offer to (Tables S31-S35). All search committee members surveyed were based at R1 universities (Box 1). A) Distribution of the fields of study and years of experience for the search committee survey respondents. B) The median number of faculty job openings, number of applicants per opening, applicants that make the first cut, applicants who are invited for phone/Skype interview and offers made. C) The quantitative rating of search committee faculty on metrics: candidate/applicant research proposal, career transition awards, postdoctoral fellowships, graduate fellowships, PI/mentor reputation (lab pedigree), Cell/Nature/Science journal publications, Impact factor of other journal publications, Teaching experience and value of preprints based on a 5-level Likert scale where 1=not at all and 5=heavily. D) Visual summary of the job applicant perception and the survey results along with the search committee survey results. A number of metrics were not measured/surveyed as part of our study. These missing values are shown in gray.

We sought to understand what factors search committees found most important, what their perception of the market was, and how they felt it had changed since they first became involved in hiring. Figure 8C shows how heavily distinct factors are weighted in making a decision as assessed from 1 (not weighted at all) to 5 (heavily weighted). We found the most important reported factors to be those most closely related to what a candidate would do going forward, such as their proposed research and current funding, rather than their previous track record (Table S35). This and other areas showed a discrepancy between faculty and applicant perceptions of metrics associated with successful candidates (Figure 8D), highlighting the importance of obtaining data such as we present here to better inform applicants.

The majority (67%) of the search committee faculty viewed preprints favorably, although their strength may not yet be equivalent to published peer-reviewed work (Figure 8, Table S33). Sixty seven percent of the search committee survey respondents mentioned that they received over 200 applicants per job posting, while 33% mentioned that their committee received 100-199 applications per cycle (Table S31). Despite these high numbers, only 5-8 applicants are typically invited to interview, with around a third (33%) of the faculty respondents noting they did not perform off-site (phone or Skype) interviews (Figure 8B; Table S31). These statistics help demonstrate the challenges that hiring committees face; the sheer volume of applicants is overwhelming, as mentioned explicitly by several search committee respondents (Table S29).

We took this opportunity to ask the search committee survey respondents if there were additional factors that they wished applicants knew when applying (Figure 9, Table S36). Several emphasized the quality of the research and papers as the most important factor for assessing prior achievement, but that a compelling and coherent research proposal is also critical and can be underdeveloped in some otherwise competitive candidates. The importance of departmental fit was also emphasized and that at the interview stage, a candidate’s assessment is in no small part predicated on interpersonal interactions with faculty members. This sentiment is also validated by a recent twitter poll citing “overall attitude/vibe” as the single most important factor for selection at the interview stage (50). Intriguingly, while one faculty respondent noted that they rarely interview any applicant without a career transition award, for instance the NIH-based “K99/R00 Pathway to Independence Award” (a situation they noted as problematic), another lamented that applicants worried too much about metrics/benchmarks anecdotally perceived to be important, such as receiving these awards. Finally, most (73%) faculty respondents noted that it was easy to identify good candidates from their submitted application, that there were too many good applicants (67%), and that candidates often underperformed at the interview stage (67%) (Figure 9, Figure S2, Table S34).

**Figure 9.**
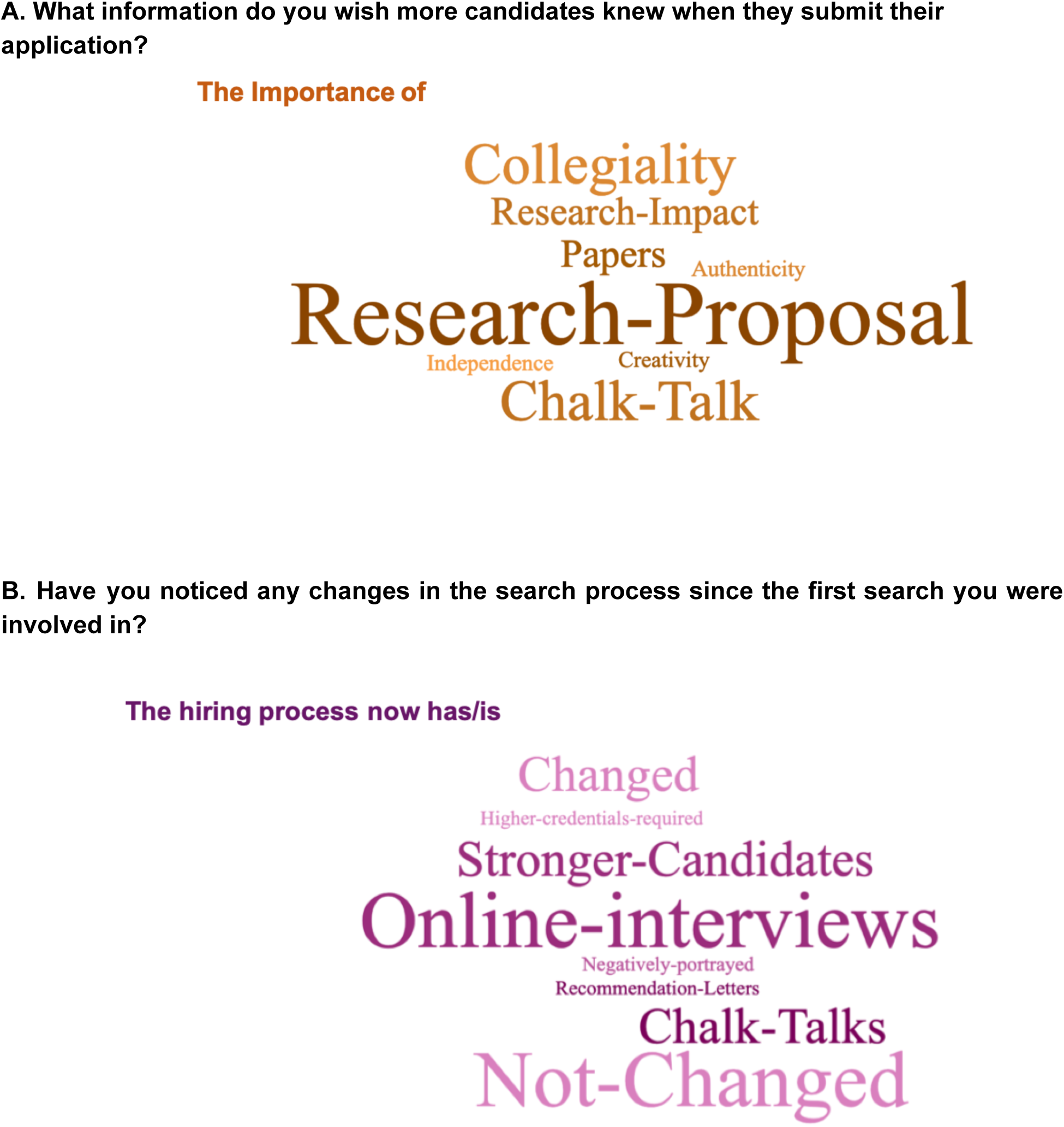
Search committee perception of the faculty job application process. Two word clouds representing responses from members of search committees in response to the following questions: A) “What information do you wish more candidates knew when they submit their application?”, and B) “Have you noticed any changes in the search process since the first search you were involved in?”. The size of the word/phrase reflects its frequency in responses, with larger phrases corresponding to more frequent responses. Search committee faculty members were able to provide long answers to both questions (Tables S36-S37).

## Discussion

### Challenges in the Academic Job Market

Currently, there is little systematic evidence for what makes a good faculty candidate. As with any opaque, high-pressure environment, an absence of clear guidelines and expectations coupled with anecdotal advice can lead individuals to focus on tangible goals and metrics that they feel will help them stand out in the system. We were able to confirm several common pieces of faculty application advice: the number of applications submitted, CNS paper first authorship, total citation count, and career transition awards (e.g. a K99/R00 award) were associated with obtaining offers. These associations held even when jointly considering all variables in a logistic regression analysis, except for the replacement of the two publication-related metrics (CNS papers and total citation count) by h-index, suggesting that they may have been proxies for “impact” in the first place. Despite these associations, these criteria were neither necessary nor sufficient for securing an offer.

CNS and other high-impact publications have been regarded as a major benchmark for trainees in the life sciences (39), and qualitative comments from our applicant survey indicated that not having a CNS paper was perceived by them to be detrimental to offer prospects. However, the majority of our respondents received offers without CNS publications, and faculty respondents on search committees did not identify as focused on CNS papers. Indeed, although applicants with a CNS 1st author paper had a higher offer percentage than those without, our logistic regression did not identify CNS 1st authorship as significant, instead detecting h-index as significant. This may suggest that CNS authors benefit from the visibility and familiarity of their work and not the title of the journal; indeed, these same advantages might be achieved by well read papers in other journals. Qualitatively, the impact of funding on research career success was well-recognized by a number of applicants and search committee members alike in our results. Quantitatively, our applicant survey found career transition awards and publication record to be among the most important factors we measured. The search committee respondents confirmed the benefit of career transition funding as major strengths for an application.

Despite challenges in the job market (2,5,24), our survey revealed positive outcomes that suggest progress in select areas. Nearly half of our applicant survey respondents reported posting at least one preprint, with several commenting that preprinting provided evidence of productivity outside of formal publication. Search committee survey respondents further confirmed that while published papers carry the most weight, preprints are generally viewed favorably. Future use of preprints as evidence of productivity records may have significant positive impacts on early career researchers, for whom timing of publications and job searches require critical considerations. Further, despite the fact that women are underrepresented at the faculty level in most STEM departments (51-54) and trail men in publication-related metrics (Fig. 2B), our data suggest very few differences in outcomes in the May 2018-May 2019 female applicant pool relative to their male counterparts. Both genders received similar numbers of interviews and offers, and female identity was even positively associated with offer status in our logistic regression analysis. In fact, these gender-based differences in publication-related metrics persisted even when considering only the 185 individuals with offers, suggesting that committees are becoming increasingly aware of gender bias in publication-related metrics and are taking them into account when evaluating applicants (Table S39).

Overall, the respondents from this survey were generally highly-qualified according to the metrics we measured, and yet they reported high stress and frustration with their experiences of the faculty job search. Applicants perceived poor mentorship as a major obstacle to their applications. Further, we found that most metrics were differentially valued by candidates and committees. Collectively, these differences in expectations between applicants and hiring institutions, coupled with the opaque requirements for obtaining a faculty position, likely drive the high stress reported by both candidates and committee members alike.

### Limitations of this study and measuring outcomes in the academic job market

There are several limitations of this study imposed by both the original survey design and general concerns, such as the anonymity of respondents, and the measurability of various contributing factors. For future data collection we suggest keeping surveys focused on region-specific job markets. Our pool of applicants were largely those seeking a position in North America. We believe these results can be aggregated, but the survey questions may not all be applicable to other large markets (e.g. Europe, China, India). We did not receive a sizable response from applicants looking outside of North America and in fields outside of life sciences to make useful comparisons. A similar survey circulated in each market individually with similar number of responses would have broader impact.

We purposely did not ask for race or ethnicity demographics, PhD or postdoc institution, and region or institution where offers were received. We believe the addition of these metrics could potentially jeopardize survey respondents’ anonymity. Despite this, these factors could be significant contributors to the receipt of an academic job offer. Racial inequalities in all STEM fields at all levels exist and need to be addressed (55), specifically with how they intersect with gender (52). The reputation of a training institution is questionably measurable, but is also often listed in anecdotal advice as important. Recently it was reported that a majority of new faculty are hired from a minority of institutions providing postdoc training (56,57). It is possible that adding institutional reputation to the other traditional metrics we measured could provide a more complete picture of the current path to a faculty position.

While we measured some of the attributes widely perceived as important in faculty hiring (e.g. funding track record), others are less easily quantified (e.g. the research proposal, lab pedigree, letters of recommendation that comments from our search committee survey revealed to be important) and data collection on these items would be highly recommended in future surveys. Addressing the quality of application materials is highly context-specific (given the field, search committee, and institutional needs) and can improve (8). Other aspects which are not directly measurable and are often cited as important for applicants in the academic job market are “fit” and “networking” (20). Applicants who responded to this survey did agree that “networking”, “conferences”, “collaborations”, and “connections” were helpful in their job search (Figure 7A). Conference organizers are also starting to include badge stickers or tags for faculty job seekers to self-identify at events. For future data collection, the number of conferences or networking events attended while applicants were on the academic job market could be measured and a relationship could be established between eventual faculty offers received and these networking metrics. Departmental or institutional “fit” is largely determined by the search committee on an individual basis. The metrics that determine a good “fit” are wildly variable and will likely never be adequately measured (58).

All questions in our survey were optional. We chose this survey design in order to make the survey easier for respondents to complete; however, missing answers represent a source of potential bias as unanswered questions may represent answers that could be negatively perceived and/or zero in value. For example, some individuals may not have felt comfortable indicating they had zero offers leading our dataset to appear overly inflated in offer percentage. Such bias could also affect the imputations in our logistic regression, and for these reasons we have attempted to provide multiple transparent and qualified analyses of the data. Future surveys may benefit from all questions requiring a response. It is also possible that participation in the survey from the outset suffers from survivorship bias, in that those applicants that had a positive experience are more likely to reflect upon it and complete a survey on the process. Our survey was also undoubtedly completed by a highly-engaged group of aspiring future faculty. The Future PI Slack group itself is a space for postdoctoral researchers most interested in obtaining a faculty career to engage with and learn from one another. Thus, the survey data likely reflects a highly motivated and accomplished group and not the full pool of applicants to faculty positions each year. Wider dissemination of future surveys will hopefully be aided by the publication of these results and increased awareness of the survey among trainees in various research communities.

Finally, the data from our job applicant survey focused on candidates and not the search committees. It is unclear how many individual searches are represented in our dataset. It is likely that as many as ∼200-500 committees were represented in our aggregated job applicant data, and different committees may adopt distinct assessment criteria. Our limited search committee survey responses show that these committees favor a holistic assessment of candidates and that decision by universal criteria (especially based solely on career transition awards or CNS publications) is a myth. Future studies may benefit from surveying a larger pool of search committees to see what major trends and practices dominate, whether the majority of searches adopt a holistic assessment approach, or if there is heterogeneity among committees in how tenure-track hiring assessments are conducted.

## Conclusions

The faculty job search process lacks transparency. This needs to change. The current system is not adequately serving the applicants, mentors, search committees, or institutions. Of over 300 responses by job applicants, we did not receive a single positive comment on the process. This is in spite of the fact that 58% of our applicants received at least one job offer. Our data suggests that baseline thresholds for who is more likely to receive a faculty job offer exist, but that there are many different paths to obtaining a job offer. The variety of paths likely reflects both the applicant’s preparation, as well as different evaluation criteria utilized by individual search committees. Increasing the transparency of the process through systematic data collection would allow a more detailed study of these many paths. It is our hope that this will not only allow all stakeholders to make informed decisions, but will also allow critical examination, discussion, and reassessment of the implicit and explicit values and biases being used to select the next generation of academic faculty. Such discussions are critical in building an academic environment that values and supports all of its members.

## Materials & Methods

### Survey Materials

We designed a survey (“applicant survey”) to collect demographics and metrics that were commonly discussed on Future PI Slack during the 2018-2019 academic job search cycle. The survey was designed to take less than 5 minutes in order to maximize response rates. Respondents were not required to answer all questions. After collecting and performing initial analyses of this survey, we designed an additional survey for search committees (“search committee survey”). The text of both surveys used in this work is included in the supplemental material as appendices. A Google form was used to conduct both surveys.

The applicant survey was distributed on various social media platforms including the Future PI Slack group, Twitter, and Facebook. The survey was also distributed by several postdoc association mailing lists including in North America, Europe and Asia. The applicant survey was open for approximately six weeks to collect responses. The search committee survey was distributed to specific network contacts of the various authors. Though this distribution was more targeted, a Google form link was still used to maintain anonymity. The search committee survey was open for approximately three weeks to collect responses. In both cases, respondents to the surveys were asked to self-report, and the information collected was not independently verified.

### Data Analysis

Microsoft Excel and RStudio were used to graph the results of both surveys shown in Figures 1-6 and 8. Specifically, data was filtered and subdivided using the “tidyverse” collection of R packages, and figure plots were generated using the “ggplot2” package. Whenever statistical analyses were used, the exact tests, p-values and **χ**^2^ values are reported in the appropriate figure or figure legend or caption, results section and Table S21, and represent the implementations in the basic R “stats” package. A p-value of less than 0.05 was considered significant. Where a number of demographics are combined in the reporting throughout this study, any analysis group with less than five respondents were combined with other similar values instead of the raw *n* value in an effort to protect the anonymity of participants. Briefly statistical methods are as follows: in general, the two-tailed Wilcoxon rank sum test (with Holm correction when applicable) or Chi-squared test was used to report p-values (see Table S21 for detailed breakdown). The qualitative survey comments were categorized by theme (keywords/context) describing each comment and the frequency of comments pertaining to a particular theme and tabulated (Tables S22, S26-27, S36-37). Word clouds were generated using the WordItOut platform (59) (Figures 7 and 9). The visual summary heatmap of the job applicant perception and the survey results along with the search committee survey results (Figure 8D) was created by counting the frequency of comments for each metric (i.e. publications, fellowships, preprints) from the qualitative (long answer) questions survey respondents (Tables S22, S26-27, S36-37). The job applicant survey quantitative results were also used to rank metrics based on significance (as determined by Wilcoxon analysis or logistic regression analysis (Table S21)) and were also incorporated into the heatmap (Figure 8D). A number of metrics were not measured/surveyed as part of our study. These missing values are shown in gray.

Logistic regression analysis was performed in R using the “glm” function with the “family” parameter set to “binomial”. All variables collected in the survey were included as independent variables, except those that were considered to be outcomes (numbers of remote interviews, onsite interviews and offers). The outcome variable was a binary “Offer” or “No offer” variable. All continuous variables were z-score normalized to ensure that they were centered and scaled consistently. When all independent variables were considered together, missing values accounted for nearly two-thirds of the data, and were therefore, imputed by fitting a bagged tree model for each variable (as a function of all the others) (60). Both variations of the analysis (missing data excluded and missing data imputed) were reported.

In order to visualize the potential paths to an offer, a decision tree was learned automatically from the data using the C5.0 algorithm (60). All possible combinations of the following parameter settings were evaluated: (1) either the tree-based variant or the rule-based variant of the algorithm was run, (2) winnowing of irrelevant variables was set to “TRUE” or “FALSE”, and (3) the number of boosting “trials” was set to 1, 4, 16, 32 or 64. The parameter combination with the best accuracy in predicting offer status in a 10-fold cross-validation experiment (as implemented in the “caret” package in R) was chosen (61). Since decision trees naturally handle missing values and differences in scales, no additional imputation or data normalization was performed before training and testing. The most accurate tree was found to be the one that used the rule-based variant, had no winnowing and no boosting (trials = 1) and was plotted using the “plot” function in the “partykit” R package (62) and then manually edited in Illustrator.

### Data availability

The authors confirm that, for approved reasons, access restrictions apply to the data underlying the findings. Raw data underlying this study cannot be made publicly available in order to safeguard participant anonymity and that of their organizations. Ethical approval for the project was granted on the basis that only aggregated data is provided (as has been provided in the supplementary tables) (with appropriate anonymization) as part of this publication.

### Statement of Ethics

This survey was created by researchers listed as authors on this publication, affiliated with universities in the United States in an effort to promote increased transparency on challenges early career researchers face during the academic job search process. The authors respect the confidentiality and anonymity of all respondents. No identifiable private information has been collected by the surveys presented in this publication. Participation in both surveys has been voluntary and the respondents could choose to stop responding to the surveys at any time. Both “Job Applicant” and “Search Committee” survey has been verified by the University of North Dakota Institutional Review Board (IRB) as Exempt according to 45CFR46.101(b)(2): Anonymous Surveys No Risk on 08/29/2019. IRB project number: IRB-201908-045. Please contact Dr. Amanda Haage for further inquiries.

## Supporting information

All supplemental figures & tables

## Acknowledgements

The authors thank Drs. Carol Greider, Feilim Mac Gabhann, Cori Bargmann, Mark Kunitomi, Needhi Bhalla, Lucia Peixoto, Sarah Stone and Dario Taraborelli for their valuable comments on an earlier version of this manuscript. This work was supported by a start up fund from University of North Dakota to AH, F32GM125388 (NIGMS) to JDF, T32HL007749 (NHLBI) to AJK, start up fund from Midwestern University to NMJ, support from the Washington Research Foundation Fund for Innovation in Data-Intensive Discovery and the Moore-Sloan Data Science Environments Project at the University of Washington to VP. The authors would like to thank the entire Future PI Slack community and those who support them in their support of this work.

