## Supplementary material for "Insights from a survey-based analysis of the academic job market": All supplemental figures & tables

### Supplementary File

Consists of 3 figures and 39 Tables referenced in the main text and draft of the 2 surveys conducted for this study.

| **Table S1.** Common online resources for finding academic jobs | |
|---|---|
| **General job search** | **Functionality** |
| https://academicjobsonline.org/ajo | Can search by discipline & location |
| https://jobs.sciencecareers.org/ | Jobs in Science & Technology from Science Careers |
| https://www.indeed.com/ | General job search from Indeed.com |
| https://www.higheredjobs.com/ | Higher Education Job search by Institution, job type, by category, school & location |
| https://www.hercjobs.org/ | Higher Education jobs by state & job type |
| https://academicpositions.com/ | Academic job search by employer |
| https://www.timeshighereducation.com/unijobs/en-us/ | Academic job search by keyword & location |
| https://chroniclevitae.com/ | Higher Education Job search by location, institution type, & field/keywords |
| https://www.interfolio.com/ | Interfolio Faculty Search is a faculty hiring platform, covering every detail of the recruitment process, from confidential letters to 100% compliance on EEO forms. |
| **Field-specific job search** | **Functionality** |
| https://jobs.ascb.org/ | Cell Biology jobs |
| http://psychjobsearch.wikidot.com/ | Psychology jobs |
| http://neurorumblr.com/ | Neuroscience jobs |
| http://chemjobber.blogspot.com/ | Chemistry jobs |
| http://ecoevojobs.net | Ecology & Evolution Biology jobs |

**Table S1.** Resources for applicants for finding academic jobs. These resources were often mentioned by our applicant survey respondents and cited by others as helpful for locating academic job ads across different fields.

| Table S2. Applicants by their Field of Research & Gender | | | | |
|---|---|---|---|---|
| **Theme** | **Total Applicant number (total n=317)** | **Female Applicant number (total n=153)** | **Male Applicant number (total n=160)** | **Non-Binary/did not disclose gender (total n=4)** |
| **Total Number of respondents to this question** | 100% (317 out of 317) | 48.3% (153 out of 317) | 50.5% (160 out of 317) | 0% (0 out of 4) |
| **Biomedical or Life Sciences** | 52.4% (166 out of 317) | 54.2% (83 out of 153) | 50.6% (81 out of 160) | 50% (2 out of 4) |
| **Biology (other)** | 19.6% (62 out of 317) | 23.5% (36 out of 153) | 16.3% (26 out of 160) | 25% (1 out of 4) |
| **Chemistry** | 8.8% (28 out of 317) | 5.9% (9 out of 153) | 11.9% (19 out of 160) | 0% (0 out of 4) |
| **Bioengineering** | 3.8% (12 out of 317) | 2.6% (4 out of 153) | 5% (8 out of 160) | 0% (0 out of 4) |
| **Earth Sciences** | 2.5% (8 out of 317) | 3.3% (5 out of 153) | 1.3% (2 out of 160) | 25% (1 out of 4) |
| **Physics** | 2.2% (7 out of 317) | 1.3% (2 out of 153) | 3.1% (5 out of 160) | 0% (0 out of 4) |
| **Chemical Engineering** | 2.2% (7 out of 317) | 2% (3 out of 153) | 2.5% (4 out of 160) | 0% (0 out of 4) |
| **Psychology** | 2.2% (7 out of 317) | 2% (3 out of 153) | 2.5% (4 out of 160) | 0% (0 out of 4) |
| **Engineering** | 1.9% (6 out of 317) | 1.3% (2 out of 153) | 2.5% (4 out of 160) | 0% (0 out of 4) |
| **Social Sciences** | 1.3% (4 out of 317) | 2.6% (4 out of 153) | 0% (0 out of 160) | 0% (0 out of 4) |
| **Computer Sciences** | 1.3% (4 out of 317) | 1.3% (2 out of 153) | 1.3% (2 out of 160) | 0% (0 out of 4) |
| **Other Fields: Health Sciences + Cognitive Sciences + Materials Science + Mathematics** | 1.9% (6 out of 317) | Less than 1% (1 out of 153) | 3.1% (5 out of 160) | 0% (0 out of 4) |

**Table S2.** Overview of job application survey respondents' (total & by gender) field of study. Fields which had fewer than 3 respondents in our job applicant survey were aggregated as "Other Fields" in the table. All percentages are calculated out of the total number of respondents.

| Table S3. Applicant Demographics: Country of Research Origin (Applicant Location) | |
|---|---|
| **Applicant Origin (country in which the applicant conducted research while applying for a faculty position)** | **Number of Applicants** |
| **United States** | 72% (214 out of 297) |
| **Canada** | 10.1% (30 out of 297) |
| **United Kingdom** | 9.1% (27 out of 297) |
| **Germany** | 2.4% (7 out of 297) |
| **France** | 1.7% (5 out of 297) |
| **Other countries: Spain + Sweden + India + Norway + Singapore + Japan + The Netherlands + Switzerland** | 4.7 % (14 out of 297) |
| **Number of applicants who indicated country of research origin (not blank)** | 93.7% (297 out of 317) |
| **Did not respond to this survey Question** | 6.3% (20 out of 317) |

**Table S3.** Overview of candidates' country of research origin. Regions which had fewer than 5 respondents in our job applicant survey were aggregated as "Other countries" in the table. All percentages are calculated out of the total number of respondents to this particular survey question (297) not total number of overall survey respondents (n=317).

| Table S4. Country to which faculty application was made (Job Location) | |
|---|---|
| **Application destination (country to which the applicants applied for a faculty position)** | **Number of Applicant** |
| **United states** | 81.7% (259 out of 317) |
| **Canada** | 33.1% (105 out of 317) |
| **United Kingdom** | 24.3% (77 out of 317) |
| **Germany** | 10.4% (33 out of 317) |
| **France** | 7.8% (24 out of 317) |

| | |
|---|---|
| **Switzerland** | 7.8% (24 out of 317) |
| **Australia** | 5 % (16 out of 317) |
| **The Netherlands** | 4.1% (13 out of 317) |
| **Austria** | 4.1% (13 out of 317) |
| **Spain** | 1.9% (6 out of 317) |
| **Sweden** | 1.9% (6 out of 317) |
| **Belgium** | 1.9% (6 out of 317) |
| **Finland** | 1.6% (5 out of 317) |
| **Other countries & regions: Taiwan + Ireland + Turkey + United Arab Emirates + China + Fiji + Grenada + Jamaica + St.Lucia + Singapore + Norway + New Zealand + Japan + Israel + Portugal + Denmark + Hong Kong** | 11% (35 out of 317) |
| **Number of applicants who responded to this survey question (not blank)** | 100% (317 out of 317) |

**Table S4.** Applicant Survey Demographics: Overview of the countries to which the faculty candidates applied to, for faculty positions. Note: most candidates applied to more than one country. Regions which had fewer than 5 respondents in our job applicant survey were aggregated as "Other countries & regions" in the table. All percentages are calculated out of the total number of respondents to this particular survey questions (n=317).

| **Table S5. Applicants' current research/academic position** | |
|---|---|
| **Theme** | **Applicant number (n=317)** |
| **Postdoctoral Researcher** | 96% (304 out of 317) |
| **Doctoral researcher (PhD)** | less than 1% (2 out of 317) |
| **Principal Investigator (PI)** | 1.3 % (4 out of 317) |
| **Other Research Position** | 2.2% (7 out of 317) |
| **Did not respond to this survey Question** | 0% (0 out of 317) |

**Table S5.** Overview of current academic position of our job applicant survey respondents. All percentages are calculated out of the total number of respondents to this particular survey questions (n=317).

| Table S6. Applicants' Postdoctoral training time statistics | |
|---|---|
| **Theme** | **Postdoctoral Training (in years)** |
| **Time spent in postdoctoral training by all** | Average = 4.35 years; Median = 4 years |
| **Time spent in postdoctoral training by Male candidates** | Average = 4.39 years; Median = 5 years |
| **Time spent in postdoctoral training by Female candidates** | Average = 4.20 years; Median = 4 years |
| **Minimum time spent in postdoctoral training by any candidates** | 1 year |
| **Maximum time spent in postdoctoral training by any candidates** | 13 years |

**Table S6.** Overview of time spent in postdoctoral training by our job applicant survey respondents.

| Table S7. Applicant Demographics: Applicants with first or multiple postdoctoral position | |
|---|---|
| **Theme** | **Number of applicants** |
| **Applicants who applied while in their first postdoctoral position (out of applicants who responded to this question)** | 68.2% (191 out of 280) |
| **Female applicants who applied while in their first postdoctoral position** | 76% (116 out of 153) |
| **Male applicants who applied while in their first postdoctoral position** | 63% (101 out of 160) |
| **Applicants who applied while in their second postdoctoral position** | 25.4% (71 out of 280) |
| **Applicants who applied while in their third postdoctoral position** | 6.4% (18 out of 280) |
| **Applicants who applied while in their 2nd-3rd postdoctoral position (out of applicants who responded to this question)** | 31.8% (89 out of 280) |
| **Responded to this survey Question** | 88.3% (280 out of 317) |
| **Did not respond to this survey Question** | 11.7% (37 out of 317) |

**Table S7.** Overview of number of postdoctoral positions that the candidates held at the time of their faculty job application. All percentages are calculated out of the total number of respondents to this particular survey questions.

| Table S8. Applicants' Scholarly metrics | | | | |
|---|---|---|---|---|
| **Theme** | **Metrics for All applicants (n=317)** | **Metrics for Female applicants (n=153)** | **Metrics for Male applicants (n=160)** | **Metrics for applicants who did not disclose gender (n=0)** |
| **Number of peer-reviewed papers** | Average=15; Median=13 | Average=13.4; Median=11 | Average=16.4; Median=14 | Average=11.3; Median=11 |
| **Number of all preprints ever posted** | Average=1.57; Median=1 | Average=0.9; Median=0 | Average=2.1; Median=1 | Average=1.6; Median=1 |
| **Number of all preprints not peer-reviewed yet** | Average=0.7; Median= 0 | Average = 0.6; Median = 0 | Average=0.8; Median= 0 | Average=1; Median=1; |
| **Number of first-authored papers** | Average=7; Median=6 | Average=6.1; Median=7 | Average=7.8; Median=5 | Average=0; Median=8 |
| **Number co-authored "CNS" papers** | Average=0.2; Median=0 | Average=0.1; Median=0 | Average=0.2; Median=0 | Average=0.3; Median=0 |
| **Number of corresponding-authored "CNS" papers** | Average=0.022; Median=0 | Average=0; Median=0 | Average=0.044; Median=0 | Average=0; Median=0 |
| **Number of all CNS papers (first author + corresponding-author + co-author)** | Average=0.405; Median=0 | Average=0.24; Median=0 | Average=0.54; Median=0 | Average=0.46; Median=0 |
| **Number of citations** | Average=426.1; Median=282 | Average=397; Median=228 | Average=455.6; Median=343 | Average=145.5; Median=145.5 |
| **Applicants who had published "CNS" papers** | 83 | 31 | 52 | 0 |
| **h-index** | Average = 8.5; Median =8 | Average = 7.9; Median =7 | Average = 9.0; Median=9 | Average = 6; Median =6 |
| **Number of Applications** | Average = 24.1; Median = 15 | Average = 21.4; Median = 13 | Average = 26.6; Median = 16.5 | Average = 30; Median = 16 |
| **Number of off-site interviews** | Average = 2.5; Median = 1 | Average = 2.9; Median = 2 | Average = 2.2; Median = 1 | Average = 2.3; Median = 1.5 |
| **Number of on-site interviews** | Average = 2.6; Median = 2 | Average = 2.8; Median = 2 | Average = 2.4; Median = 2 | Average = 3.8; Median = 2 |
| **Number of all offers** | Average = 1.1; Median = 1 | Average = 1.3; Median = 1 | Average = 1; Median = 1 | Average = 2.5; Median = 0.5 |

| | | | | |
|---|---|---|---|---|
| **Number of applicants with One or more offers** | 58.3% (183 out of 317) | 60% (92 out of 153) | 56.9% (91 out of 160) | 50% (2 out of 4) |
| **Number of NO offers** | 41.6% (132 out of 317) | 40% (61 out of 153) | 43.1% (69 out of 160) | 50% (2 out of 4) |
| **Number of applicants with One or more on-site interviews** | 77.6% (246 out of 317) | 75.8% (116 out of 153) | 79.4% (127 out of 160) | 75% (3 out of 4) |
| **Number of NO on-site interviews** | 22.4% (71 out of 317) | 24.1% (37 out of 153) | 20.6% (33 out of 160) | 25% (1 out of 4) |
| **Number of applicants with One or more off-site interviews** | 70% (220 out of 317) | 71.9% (110 out of 153) | 68.1% (109 out of 160) | 75% (3 out of 4) |
| **Number of NO off-site interviews** | 30% (95 out of 317) | 28.1% (43 out of 153) | 31.9% (51 out of 160) | 25% (1 out of 4) |

**Table S8.** Overview of the job applicant publication metrics (average citation number, average h-index, average number of peer-reviewed papers, average number of preprints, average number of peer-reviewed first-author papers, number of Cell/Nature/Science journal publications or “CNS” papers of any type meaning 1st author, co-author or corresponding author) of our survey respondents by gender breakdown.

**Table S9. Applicant Fellowship type Funding Record**

| **Theme** | **Pre-doctoral Fellowship** | **Post-doctoral Fellowship** | **Both types of fellowships** | **Received any type of fellowship** | **Received None** | **Did not Respond to this question** |
|---|---|---|---|---|---|---|
| **All Applicants (n=317)** | 14.9% (47 out of 315) | 26.7% (84 out of 315) | 37.8% (119 out of 315) | 80% (252 out of 315) | 20% (63 out of 315) | less than 1% (2 out of 317) |
| **Applicants who applied to PUIs** | 19.2% (20 out of 104) | 28.8% (30 out of 104) | 26.9% (28 out of 104) | 77.9% (81 out of 104) | 22.1% (23 out of 104) | ~1% (1 out of 104) |
| **Female Applicants (n=153)** | 15.8% (24 out of 152) | 23% (35 out of 152) | 48.7% (74 out of 152) | 87.5% (133 out of 152) | 12.5% (19 out of 152) | less than 1% (1 out of 152) |
| **Female applicants who** | 15.5% (9 out of 58) | 27.6% (16 out of 58) | 37.9% (22 out of | 82.8% (48 out of | 17.2% (10 out of | 1.7% (1 out of |

| | | | | | | |
|---|---|---|---|---|---|---|
| **applied to PUIs** | | | 58) | 58) | 58) | 58) |
| **Male Applicants (n=160)** | 13.8% (22 out of 159) | 30.2% (48 out of 159) | 27.7% (44 out of 159) | 72.3% (115 out of 159) | 27.7% (44 out of 159) | less than 1% (1 out of 159) |
| **Male applicants who applied to PUIs** | 25% (11 out of 44) | 31.8% (14 out of 44) | 13.6% (6 out of 44) | 70.5% (31 out of 44) | 29.5% (13 out of 44) | 0% (0 out of 44) |
| **Non-Binary Applicants (0)** | 0 | 0 | 0 | 0 | 0 | 0 |
| **Preferred not to disclose gender(n=4)** | 25% (1 out of 4) | 25% (1 out of 4) | 25% (1 out of 4) | 0 | 0 | 0 |
| **Preferred not to disclose gender who applied to PUIs** | 50% (1 out of 2) | 0% (0 out of 2) | 0% (0 out of 2) | 100% (2 out of 2) | 0% (0 out of 2) | 50% (1 out of 2) |
| **Theme** | Responded PhD+Both PhD & Postdoc Fellowships | | Responded Postdoc+Both PhD & Postdoc Fellowships | | | |
| **Female Applicants** | 65% (98 out of 152) | | 71.7% (109 out of 152) | | | |
| **Male Applicants** | 41.5% (66 out of 159) | | 57.9% (92 out of 159) | | | |
| **Preferred not to disclose gender** | 50% (2 out of 4) | | 50% (2 out of 4) | | | |

**Table S9.** Overview of the types of funding held by our job applicant survey respondents. Percentages are calculated out of the total number of respondents to this particular survey questions. All percentages are calculated out of the total number of respondents to this particular survey questions. Our survey questions did not distinguish between the types (e.g. government funded vs privately funded, full vs partial salary support) or number of fellowships applied to; many of these factors are likely critical in better understanding gender differences in fellowship support.

| Table S10. Applicant record of Career transition awards | | | | | |
|---|---|---|---|---|---|
| **Theme** | **Postdoctoral-to-faculty transition award of any kind the(e.g. K99, K01)** | **Co-PI grants** | **Total applicants who held a transition award or a co-PI grant** | **None (No independent funding)** | **Did not Respond to this question** |
| **All Applicants (n=317)** | 15.9% (50 out of 314) | 10.5% (33 out of 314) | 25.1% (79 out of 314) | 74.8% (235 out of 314) | Less than 1% (3 out of 317) |
| **Female Applicants (n= 153)** | 17.1% (26 out of 152) | 12.5% (19 out of 152) | 28.3% (43 out of 152) | 71.7% (109 out of 152) | Less than 1% (1 out of 153) |
| **Male Applicants (n=160)** | 15.2% (24 out of 158) | 8.2% (13 out of 158) | 20.9% (33 out of 158) | 79.1% (125 out of 158) | 1.2% (2 out of 160) |
| **Non-Binary Applicants (n=0)** | 0 | 0 | 0 | 0 | 0 |
| **Preferred not to disclose gender (n=4)** | 0 | 25% (1 out of 4) | 75% (3 out of 4) | 25% (1 out of 4) | 0 |

**Table S10.** Overview of the types of transition/independent type funding held by our faculty candidate (applicant survey) respondents. Percentages are calculated out of the total number of respondents to this particular survey questions. Being a "Co-PI" of a grant as a postdoctoral researcher or research scientist means co-writing a grant with a PI (an independent investigator). The co-writer may or may not be explicitly mentioned on the grant as a Co-PI.

| Table S11. Application statistics | | | | |
|---|---|---|---|---|
| **Theme** | **Total Applicant number (n=317)** | **Female Applicants responded to this Q (n=131)** | **Male Applicants (n=160)** | **Did not disclose gender (n=4)** |
| **Total Number of all Applications made** | 7,644 | 3,268 | 4,256 | 120 |
| **Total Number of off-site interviews** | 805 | 437 | 359 | 9 |

| | | | | |
|---|---|---|---|---|
| **Total Number of on-site interviews** | 832 | 431 | 386 | 15 |
| **Total Number of offers** | 359 | 196 | 153 | 10 |
| **Approximate Number of rejections** | 2,920 | 1,118 | 1,742 | 60 |
| **Total number of No Feedbacks (did not hear anything back)** | 4,365 | 2,150 | 2,514 | 60 |
| **Did not respond to this survey Question** | 0 | 22 | 0 | 0 |
| **Mean Number of Applications made** | Average = 24.1; Median = 15 | Average = 21.4; Median = 13 | Average = 26.6; Median = 16.5 | Average = 30; Median = 16 |
| **Number of applicants with at least one off-site interviews** | 70% (220 out of 317) | 71.9% (110 out of 153) | 68.1% (109 out of 160) | 75% (3 out of 4) |
| **Number of applicants with at least one on-site interviews** | 77.6% (246 out of 317) | 75.8% (116 out of 153) | 79.4% (127 out of 160) | 75% (3 out of 4) |
| **Number of applicants with at least one offer** | 58.3% (185 out of 317) | 60% (92 out of 153) | 56.9% (91 out of 160) | 50% (2 out of 4) |

**Table S11.** Overview of application statistics: total number of applications made, offsite (remote via phone or online via Skype) interviews, onsite interviews, offers made, approximate number of rejections and total number of no feedbacks received from faculty job committees to our survey respondents.

| **Table S12. Twitter Poll #1: Number of offers current faculty received** | | | | |
|---|---|---|---|---|
| **Theme** | **1** | **2-3** | **4+** | **Just show me the poll results** |
| **Number of Respondents (n=749)** | 21% (167) | 20% (160) | 8% (63) | 51% (404) |

**Table S12.** Overview of the responses to a twitter poll with the question: “Faculty, when you accepted your first position, how many offers did you have to chose from?”.

| Table S13. Applicants who also applied to *Non-faculty jobs* | |
|---|---|
| **Theme** | **Applicant number** |
| **Total Number of Applicants who responded to this question** | 99.4% (315 out of 317) |
| **Yes applied to non-faculty jobs** | 28.6% (90 out of 315) |
| **Research scientist in federal government** | 1.3% (4 out of 315) |
| **R&D positions in biotech companies** | 1.6% (5 out of 315) |
| **Non-Governmental Organizations (NGO)** | Less than 1% (1 out of 315) |
| **National Institutes of Health (NIH)** | Less than 1% (1 out of 315) |
| **Science communication job** | Less than 1% (2 out of 315) |
| **Community involvement job** | Less than 1% (1 out of 315) |
| **Non-profit organizations** | Less than 1% (1 out of 315) |
| **Museum Curation (non-profit, government)** | Less than 1% (2 out of 315) |
| **Geophysical Laboratory of a research Institution of Science(Non-profit)** | Less than 1% (1 out of 315) |
| **Private laboratories** | Less than 1% (1 out of 315) |
| **Zoos** | Less than 1% (1 out of 315) |
| **No did NOT apply to non-faculty jobs** | 71.4% (225 out of 315) |
| **Did not respond to this survey Question** | Less than 1% (2 out of 317) |

**Table S13.** Overview of candidates who also applied for non-faculty jobs (e.g. Industry positions, government jobs, etc). Percentages are calculated out of the total number of respondents to this particular survey questions (n=315 applicants).

| Table S14. Applicant responses on Cell/Nature/Science or "CNS" journal publications | | | | |
|---|---|---|---|---|
| **Theme** | **Metrics for All applicants (n=317)** | **Metrics for Female applicants (n=153)** | **Metrics for Male applicants (n=160)** | **Metrics for applicants who did not disclose gender (n=4)** |
| **Number of applicants who responded to this survey question who** | 26% (81 out of 311) | 20.1% (30 out of 149) | 32.1% (50 out of 156) | 33.3% (1 out of 3) |

| | | | | |
|---|---|---|---|---|
| **had published at least one "CNS" papers** | | | | |
| **Published One "CNS" paper of any authorship type out of those applicants who published CNS** | 77.8% (63 out of 81) | 86.7% (26 out of 30) | 74% (37 out of 50) | 0% (0 out of 1) |
| **Published any 1st author "CNS" paper out all applicants** | 16.1% (50 out of 311) | 5.8% (18 out of 311) | 10% (31 out of 311) | Less than 1% (1 out of 311) |
| **Published more than one "CNS" paper of any authorship type** | 23.4% (19 out of 81) | 13.3% (4 out of 30) | 28% (14 out of 50) | 100% (1 out of 1) |
| **Number of applicants who responded to this survey question and had Not published "CNS" papers** | 73.3% (228 out of 311) | 79.9% (119 out of 149) | 67.9% (106 out of 156) | 66.7% (2 out of 3) |
| **Did not respond to this survey Question** | 2.5% (8 out of 317) | 2.6% (4 out of 153) | 2.5% (4 out of 160) | 25% (1 out of 4) |
| **Applicants who published One 1st author "CNS" paper** | 51.8% (42 out of 81) | 53.3% (16 out of 30) | 52% (26 out of 50) | 0% (0 out of 1) |
| **Applicants who published Two 1st author "CNS" papers** | 9.9% (8 out of 81) | 6.7% (2 out of 30) | 10% (5 out of 50) | 100% (1 out of 1) |
| **Applicants who published One co-author "CNS" paper** | 42% (34 out of 81) | 50% (15 out of 30) | 36% (18 out of 50) | 100% (1 out of 1) |
| **Applicants who published more than one co-author "CNS" paper** | 7.4% (6 out of 81) | 3.3% (1 out of 30) | 10% (5 out of 50) | 0% (0 out of 1) |
| **Applicants who published One corresponding author "CNS" paper** | 2.5% (2 out of 81) | 0% (0 out of 30) | 4% (2 out of 50) | 0% (0 out of 1) |
| **Applicants who published more than** | 2.5% (2 out of 81) | 0% (0 out of 30) | 4% (2 out of 50) | 0% (0 out of 1) |

| |
|---|
| **one corresponding author "CNS" paper** |

**Table S14.** Overview of the number of Cell/Nature/Science ("CNS") journal publications of our job applicant survey respondents by gender breakdown. Percentages are calculated out of the total number of respondents to this particular survey questions.

**Table S15. Applicant responses to the question on number of Active Preprints (all preprints online not peer-reviewed yet) at the time of application**

| Theme | Applicant number (All n=317) | Female Applicants (n=153) | Male Applicants (n=160) | Did not disclose gender applicants (n=4) |
|---|---|---|---|---|
| **Applicants who responded to this survey Question** | 94% (298 out of 317) | 45.4% (144 out of 317) | 47.6% (151 out of 317) | Less than 1% (3 out of 317) |
| **Yes Had posted unpublished Preprints** | 39.6% (118 out of 298) | 36.8% (53 out of 144) | 41.7% (63 out of 151) | 66.7% (2 out of 3) |
| **Posted 1 unpublished preprint** | 21.5% (64 out of 298) | 20.8% (30 out of 144) | 21.9% (33 out of 151) | 33.3% (1 out of 3) |
| **Posted 2 unpublished preprints** | 12.1% (36 out of 298) | 10.4% (15 out of 144) | 13.2% (20 out of 151) | 33.3% (1 out of 3) |
| **Posted 3 unpublished preprints** | 2.7% (8 out of 298) | 4.2% (6 out of 144) | 1.3% (2 out of 151) | 0% (0 out of 3) |
| **Posted more than 3 unpublished preprints** | 3.4% (10 out of 298) | 1.4% (2 out of 144) | 5.3% (8 out of 151) | 0% (0 out of 3) |
| **No Had NOT posted unpublished Preprints** | 60.4% (180 out of 298) | 63.2% (91 out of 144) | 58.3% (88 out of 151) | 33.3% (1 out of 3) |
| **Did not respond to this survey Question** | 6.0% (19 out of 317) | 5.9% (9 out of 153) | 5.6% (9 out of 160) | 25% (1 out of 4) |

| Total number of all Career Preprints published by Applicants | | | | |
|---|---|---|---|---|
| **Theme** | **Applicant number (All n=317)** | **Female Applicants (n=153)** | **Male Applicants (n=160)** | **Did not disclose gender applicants (n=4)** |
| **Applicants who responded to this survey Question** | 85.1% (270 out of 317) | 39.4% (125 out of 317) | 44.8% (142 out of 317) | Less than 1% (3 out of 317) |
| **Yes Had posted Preprints throughout career** | 54.8% (148 out of 270) | 48% (60 out of 125) | 60.6% (86 out of 142) | 66.7% (2 out of 3) |
| **Posted 1 preprint** | 20% (54 out of 270) | 24% (30 out of 125) | 16.2% (23 out of 142) | 33.3% (1 out of 3) |
| **Posted 2 preprints** | 14.8% (40 out of 270) | 13.6% (17 out of 125) | 16.2% (23 out of 142) | 0% (0 out of 3) |
| **Posted 3 preprints** | 6.3% (17 out of 270) | 4.8% (6 out of 125) | 7.7% (11 out of 142) | 0%(0 out of 3) |
| **Posted more than 3 preprints** | 13.7% (37 out of 270) | 5.6% (7 out of 125) | 20.4% (29 out of 142) | 33.3%(1 out of 3) |
| **No Had NOT posted Any Preprints** | 45.2% (122 out of 270) | 52% (65 out of 125) | 39.4% (56 out of 142) | 33.3%(1 out of 3) |
| **Did not respond to this survey Question** | 13.9% (47 out of 317) | 18.3% (28 out of 153) | 11.3% (18 out of 160) | 25%(1 out of 4) |

**Table S15.** Overview of candidates who had unpublished preprints at the time of their job application. Percentages are calculated out of the total number of respondents to this particular survey question.

| Table S16. Applicant responses on Patenting of their research | | | | |
|---|---|---|---|---|
| **Theme** | **Applicant number** | **Female Applicants (n=153)** | **Male Applicants (n=160)** | **Did not disclose gender (n=4)** |
| **Total Number of Applicants who responded to this survey question** | 94%<br>(298 out of 317) | 94.1%<br>(144 out of 153) | 94.4%<br>(151 out of 160) | 75%<br>(3 out of 4) |
| **Yes Had approved or pending patents** | 18.8%<br>(56 out of 298) | 15.3%<br>(22 out of 144) | 22.5%<br>(34 out of 151) | 0%<br>(0 out of 3) |
| **had 1 approved or pending patent** | 12.1%<br>(36 out of 298) | 11.8%<br>(17 out of 144) | 12.6%<br>(19 out of 151) | 0%<br>(0 out of 3) |
| **had 2 approved or pending patents** | 4%<br>(12 out of 298) | 2.1%<br>(3 out of 144) | 6%<br>(9 out of 151) | 0%<br>(0 out of 3) |
| **had 3 approved or pending patents** | less than 1%<br>(2 out of 298) | 1.4%<br>(2 out of 144) | 0%<br>(0 out of 151) | 0%<br>(0 out of 3) |
| **had more than 3 approved or pending patents** | 2%<br>(6 out of 298) | 0%<br>(0 out of 144) | 4%<br>(6 out of 151) | 0%<br>(0 out of 3) |
| **No did NOT hold any pending or approved patents** | 81.2%<br>(241 out of 298) | 84.7%<br>(122 out of 144) | 77.5%<br>(117 out of 151) | 100%<br>(3 out of 3) |
| **Preferred Not to disclose information** | 6%<br>(19 out of 317) | 5.9%<br>(9 out of 153) | 5.6%<br>(9 out of 160) | 25%<br>(1 out of 4) |

**Table S16.** Overview of Candidates who had approved or pending patents from their research at the time of their job application. Percentages are calculated out of the total number of respondents to this particular survey questions.

| Table S17. Applicants by their application type (R1 Universities, PUIs or both) & Gender | | | | |
|---|---|---|---|---|
| **Theme** | **Total Applicant number** | **Female Applicant number** | **Male Applicant number** | **Did not disclose gender Applicant number** |
| **Total Number of Applicants** | 100%<br>(317 out of 317) | 48.3%<br>(153 out of 317) | 50.5%<br>(160 out of 317) | 1.3%<br>(4 out of 317) |
| **R1 universities** | 67.2%<br>(213 out of 317) | 62.7%<br>(96 out of 153) | 72.5%<br>(116 out of 160) | 50%<br>(2 out of 4) |

| | | | | |
|---|---|---|---|---|
| **PUIs only** | 7.9%<br>(25 out of 317) | 9.8%<br>(15 out of 153) | 6.3%<br>(10 out of 160) | 0%<br>(0 out of 4) |
| **Both R1 & PUIs** | 24.9%<br>(79 out of 317) | 13.6%<br>(43 out of 317) | 11%<br>(34 out of 317) | 50%<br>(2 out of 4) |
| **Did not respond to this survey Question** | 0%<br>(0 out of 317) | 0%<br>(0 out of 153) | 0%<br>(0 out of 160) | 0%<br>(0 out of 4) |
| **PUI Applicants' First Author paper record** | | | | |
| **Total Number of Applicants** | **Total Applicant number** | **Female Applicant number** | **Male Applicant number** | **Did not disclose gender Applicant number** |
| **Applicants who applied to PUIs only** | Max = 9;<br>Average = 4;<br>Median = 4;<br>Min = 1 | Max = 7;<br>Average = 3.4;<br>Median = 4;<br>Min = 1 | Max = 9;<br>Average = 4.2;<br>Median = 4;<br>Min = 1 | Max = 0 ;<br>Average = 0;<br>Median = 0;<br>Min = 0 |
| **Applicants who applied to Both R1 & PUIs (PUI only + both responses)** | Max = 25;<br>Average = 6.3 ;<br>Median = 6;<br>Min = 1 | Max = 20;<br>Average = 5.6;<br>Median = 5;<br>Min = 1 | Max = 25;<br>Average = 7.3;<br>Median = 6.5;<br>Min = 1 | Max = 5;<br>Average = 5;<br>Median = 5;<br>Min = 5 |
| **Applicants who applied to PUIs & both R1 + PUI & Did not respond to this survey Question on their 1st author papers** | 0%<br>(0 out of 104) | 0%<br>(0 out of 58) | 0%<br>(0 out of 44) | 50%<br>(1 out of 2) |

**Table S17** Overview of job application survey respondents' (total & by gender) applications to R1 Universities (high-activity Research Universities), PUIs (Primarily Undergraduate Institutions) (see Box1 for definitions) or applied to both types of institutions. Percentages are calculated out of the total number of respondents to this particular survey questions.

| **Table S18. Applicant Teaching experience** | | | |
|---|---|---|---|
| **Theme** | **No teaching experience** | **Yes-Teaching Assistantship (TA)** | **Yes-Beyond TA** |
| **All Applicants (n=317)** | ~1% (3 out of 317) | 45.1%<br>(143 out of 317) | 53.9%<br>(171 out of 317) |

| | | | |
|---|---|---|---|
| **Applicants who applied to PUIs** | 0%<br>(0 out of 102) | 42.2%<br>(43 out of 102) | 57.8%<br>(59 out of 102) |
| **Female Applicants(n=153)** | Less than 1%<br>(1 out of 153) | 42.5%<br>(65 out of 153) | 56.9%<br>(87 out of 153) |
| **Female Applicants who applied to PUIs** | 0% (0 out of 58) | 43.1%<br>(25 out of 58) | 56.9%<br>(33 out of 58) |
| **Male Applicants(n=160)** | 1.3%<br>(2 out of 160) | 48.8%<br>(78 out of 160) | 50%<br>(80 out of 160) |
| **Male Applicants who applied to PUIs** | 0%<br>(0 out of 44) | 40.9%<br>(18 out of 44) | 59.1%<br>(26 out of 44) |
| **Non-Binary Applicants(n=0)** | 0 | 0 | 0 |
| **Preferred not to disclose gender(n=4)** | 0 | 0 | 1.3%<br>(4 out of 317) |

**Table S18.** Overview of the teaching experience (Teaching Assistantship for a course (lecture based and/or laboratory based) for the course instructor only versus beyond teaching assistantship which is independently designing and instructing undergraduate and/or graduate courses) of our applicant survey respondents. Percentages are calculated out of the total number of respondents to this particular survey questions.

**Table S19. Themes from applicant written responses to specific types of teaching experiences they had beyond teaching assistantship**

| **Theme** | **Example Survey Responses** | **Frequency of the response *n*** |
|---|---|---|
| **Instructed Undergraduate Courses** | Taught undergraduate course as instructor of record | (55) |
| **Co-Instructed Undergraduate Courses** | I have been a primary or co-instructor for several undergraduate classes | (10) |
| **Guest lectured Undergraduate courses** | Guest lecturing undergraduates, mentoring undergraduates. | (8) |
| **Instructed Graduate Courses** | Lectured medical school graduate school ~20 contact hours in first year, large multi-lecturer courses | (28) |

| | | |
|---|---|---|
| **Co-Instructed Graduate Courses** | co-taught course as a software and data carpentry instructor | (7) |
| **Guest-Lectured Graduate Courses** | Guest lecturing undergraduates, mentoring undergraduates | (5) |
| **Independent Instructor/Lecturer (type of course not specified)** | I taught two 1-year lectures. | (22) |
| **Co-Instructor/Lecturer (type of course not specified)** | Lecture an undergrad stats class for 2 years as an invited lecturer | (2) |
| **Guest-Lecturer (type of course not specified)** | Multiple guest lecturing occasions | (14) |
| **Lab Course Instructor** | Laboratory instructor for 5 years,3 years as Lab course teacher as a postdoctoral fellow. | (2) |
| **Lecturing for Workshops** | leading domain workshops at my university and others | (8) |
| **Instructor for High School Courses** | High school teacher for 1 year, taught college level summer courses for high school students. | (2) |
| **Visiting Assistant Professorship** | Visiting Professor at a liberal arts college | (2) |
| **Adjunct Teaching Instructor for Undergraduate Courses at a Community College or PUI** | Adjunct faculty for one year, taught two undergraduate courses as faculty of record and prepared all the materials for both courses. | (16) |
| **Adjunct Teaching Instructor for Undergraduate courses at an R1 University** | Adjunct faculty, designed graduate course for 1 semester. | (2) |
| **Total Adjunct teaching positions (i.e. all college level teaching counts)** | College level teaching experience (Adjunct undergraduate instructor or lecturer) | (29) |
| **Teaching Certificate** | Graduate student teaching certificate | (4) |
| **Teaching Assistant for Undergraduate or Graduate Courses** | 1 semester as TA as a graduate student. | (2) |

**Table S19.** Overview of specific types of teaching experience of our job applicant survey respondents detailed in a comment question. The "Adjunct Teaching Instructor for Undergraduate Courses at a Community College or PUI" and "Adjunct Teaching Instructor for Undergraduate Courses at an R1 or PU Institution" were explicitly mentioned comments by our applicant survey respondents. The "Total Adjunct teaching positions" were the total head-count of "adjunct type" college teaching performed by our job applicant survey respondents. A total of n=162 applicants responded to this comment type long answer question.

| **Table S20. Applicants' use of resources that offered information about the application process** | | | |
|---|---|---|---|
| **Got help from** | **Said Yes Used & Yes Useful** | **Number of applicants who received at least Offers** | **Said No** |
| **Future PI Slack** | 9.5% & 8.5%<br>(30 out of 317 & 27 out of 317) | 66.7%<br>(20 out of 30) | 10.7%<br>(34 out of 317) |
| **Chemblogger** | 1.6%<br>(5 out of 317) | 40%<br>(2 out of 5) | - |
| **EcoEvoJobsWiki** | 1.3%<br>(4 out of 317) | 75%<br>(3 out of 4) | - |
| **PsychJobsWiki** | Less than 1%<br>(1 out of 317) | 100%<br>(1 out of 1) | - |

**Table S20.** Overview of candidates who were familiar with the Future PI Slack resource and other resources during their application process. Responses to "Did you find the Future PI google sheet/Slack helpful? Yes/No" Survey participants were able to provide long answer to this comment question (Future PI Slack or FPI Slack is a slack group comprised of postdoctoral researchers aspiring to apply for faculty positions).

| **Table S21. Summary of the Statistical Analysis in this paper** | | | |
|---|---|---|---|
| **Context** | **p-value** | **Statistical test used** | **Corresponding section, Figure & Table** |
| Demographics of our applicant survey respondents: years of postdoctoral training | Life sciences vs Other fields<br>p-value = $6.5 \times 10^{-6}$,<br><br>Years as Postdoc Male vs Female | Two-tailed Wilcoxon rank sum test | Results,<br>Figure 1D,<br>Table S6 |

| | | | |
|---|---|---|---|
| | Applicants<br>p-value=0.2(ns) | | |
| Number of applicant first-author papers by gender | p-value = $1.4 \times 10^{-4}$ | Two-tailed Wilcoxon rank sum test | Results, Figure 2B, Table S8 |
| Total Publications by gender | p-value = $3 \times 10^{-3}$ | Two-tailed Wilcoxon rank sum test | Results, Figure 2B, Table S8 |
| All Citations by gender | p-value = $1.5 \times 10^{-2}$ | Two-tailed Wilcoxon rank sum test | Results, Figure 2B, Table S8 |
| Applicant h-index by gender | p-value = $5.4 \times 10^{-3}$ | Two-tailed Wilcoxon rank sum test | Results, Figure 2B, Table S8 |
| Number of first-author CNS papers | Life sciences versus Other fields<br>p-value= $1.23 \times 10^{-2}$,<br><br>Male vs Female Applicants<br>p-value= $4.54 \times 10^{-2}$ | Two-tailed Wilcoxon rank sum test | Results, Figure 2C, Table S14 |
| Percentage of PhD & Postdoctoral Fellowships received by gender | p-value = $2.4 \times 10^{-3}$ | Chi-squared test<br>( $\chi^2$ =12.10, df=2) | Results, Figure 2D, Table S9 |
| Number of off-site interviews by gender | p-value = $4.1 \times 10^{-24}$ | Two-tailed Wilcoxon rank sum test | Results, Figure 3D, Table S8 |
| Applications by gender<br><br>The median number of offers also did not vary by gender. | p = $7.25 \times 10^{-2}$,<br>off-sites p = 0.1479,<br>on-sites p = 0.5813<br><br>p = 0.1775 | Wilcoxon rank-sum test<br><br>Wilcoxon rank-sum test | Results |
| Number of on-site interviews by gender | p-value = $1.2 \times 10^{-13}$ | Two-tailed Wilcoxon rank sum test | Results, Figure 3D, Table S8 |
| Number of offers that applicants received by gender | p-value = $5.0 \times 10^{-5}$ | Two-tailed Wilcoxon rank sum test | Results, Figure 3D, |

| | | | |
|---|---|---|---|
| | | | Table S8 |
| Percentage of applicants that applied for faculty jobs vs applicants that applied to other jobs | p-value = $1.9 \times 10^{-3}$ | Two-tailed Wilcoxon rank sum test | Results, Figure 3E, Table S13 |
| Percentage of applicants that had CNS publications and received off-site, on-site interviews or offers | Off-site interviews p-value = 0.33,<br><br>On-site interviews p-value = $2.7 \times 10^{-4}$,<br><br>Offers p-value = $1.5 \times 10^{-4}$ | Two-tailed Wilcoxon rank sum test | Results, Figure 4A, Table S14 |
| Correlational Analysis of scholarly metrics & offers received | CNS 1st authors Papers p-value = $1.7 \times 10^{-3}$,<br><br>Independent Funding p-value = $2.5 \times 10^{-2}$,<br><br>Total Citations p-value = $2.92 \times 10^{-2}$,<br><br>Years on the Job Market p-value = $3.45 \times 10^{-2}$,<br><br>Postdoc Fellowship p-value = 0.1690,<br><br>Total Publications p-value = 0.1904,<br><br>CNS co-authorship papers p-value = 0.3353,<br><br>H-index p-value = 0.5724,<br><br>Years as Postdoc p-value = 1.0,<br><br>1st Author papers p-value = 1.0,<br><br>Patents p-value = 1.0, | Two-tailed Wilcoxon rank sum test with Holm correction | Results, Figure 4B, Tables S6, S9, S14, S15, S16 |

| | | | |
|---|---|---|---|
| | PhD Fellowship<br>p-value = 1.0 | | |
| Breakdown of respondents by significant criteria: citation count, independent funding, and offers<br><br>Pie-charts: Breakdown of respondents with independent funding into groups with/without CNS papers and offers | citation count (all publications) distributions for groups by CNS 1st Authorship<br>p-value = $4 \times 10^{-2}$<br><br>independent funding, and offers<br>p-value = $4 \times 10^{-2}$<br><br>Applicants with CNS<br>p-value = 0.5587<br><br>Applicants without CNS<br>p-value = 0.166 | Two-tailed Wilcoxon rank sum test<br><br>Chi-squared test<br>($\chi^2$ = 0.34188, df = 1),<br><br>Chi-squared test<br>($\chi^2$ =1.9183, df = 1), | Results, Figure 4C |
| Teaching experience & R1/PUI type application | p-value = 0.5592 | Chi-squared test<br>($\chi^2$ = 0.410) | Results, Figure 5C, Table S18 |
| Teaching experience & offer percentage | p-value = 0.1633 | Two-tailed Wilcoxon rank sum test | Results, Figure 5D, Table S18 |
| Percentage of 1st authored papers by PUI applicants by gender | p-value = 0.8882 | Two-tailed Wilcoxon rank sum test | Results, Figure 6B, Table S18 |
| Percentage of Applicants that had adjunct lectureship teaching experience | p-value = $4.997 \times 10^{-4}$ | Chi-squared test<br>($\chi^2$ = 27.515) | Results, Figure 6F, Table S19 |
| Applicants who applied to PUI only or R1 or both by adjunct teaching experience | PUI only<br>p-value=0.5538,<br><br>R1 or Both R1 & PUI<br>p-value=0.9896 | Two-tailed Wilcoxon rank sum test | Results, Figure 6G |
| Median numbers by gender: application, remote(off-site) & on-site interviews, offer of a faculty position | p-values:<br>gender=0.0725,<br>off-site=0.1479,<br>on-site=0.5813, | Two-tailed Wilcoxon rank sum test | Results, Page 10 |

| | | | |
|---|---|---|---|
| | offer=0.1775 | | |
| CNS publications with at least one offer | p-value=0.033 | Chi-squared test ($\chi^2$ = 4.4871) | Results |

**Table S21.** Summary of statistical analysis. In this table and relevant figures, **"ns"** stands for not significant.

| **Table S22. Themes from Applicant survey written responses to what general perception of the application process was** | | |
|---|---|---|
| **Theme** | **Survey Responses** | **Frequency of the response *n*** |
| **Lack of Feedback** | Very little feedback about why my application was unsuccessful. We never get feedback on our application material, so it is very hard to understand how to improve. | (13) |
| **Lack of Mentorship** | I didn't really have a mentorship on how to handle the on-site interview and structure my proposed work talk. | (1) |
| **Biased** | I feel that the process is very biased and racist. | (1) |
| **Unhealthy** | I thought the application process was incredibly stressful to the point of it being unhealthy. | (1) |
| **Despair** | I found the process very frustrating. | (1) |
| **Time-Consuming** | Ultimately, the process did take an enormous amount of time and effort and caused me a great deal of anxiety. | (2) |
| **Lack of a single centralized system** | I would love to have a single centralized system, where you upload a single application and choose a (limited) number of schools to apply to. It was also painful because many schools use different application portals with different requirements. | (2) |
| **Difficult** | My health has significantly deteriorated from this process. | (3) |
| **Awful** | The process was awful. | (1) |
| **Futile** | The application process was an awful exercise in futility. | (2) |
| **System not working** | It makes you feel like the system is not working, that something has to change. | (1) |

| | | |
|---|---|---|
| **Tough** | The process is tough. | (1) |
| **Painful** | It was also painful because many schools use different application portals with different requirements. | (3) |
| **Stressful** | It is a ridiculously stressful process. I never realized that so many people have similar anxieties, insecurities, and questions as me. | (4) |
| **Depressing** | I found the process very depressing. | (1) |
| **Arduous** | The application process was hard. | (1) |
| **Demoralizing** | I really enjoyed the one interview that I had but otherwise I found the process fairly demoralizing. | (1) |
| **Terrible** | The whole process is terrible.<br>The Skype interviews are terrible. | (2) |
| **Demeaning** | The Skype interviews are demeaning, and the poor connections involved seem designed to make understanding difficult. I had one Skype interview in which the point was for the selection committee to get to laugh at me and ridicule my work. | (1) |
| **Isolating** | Applying for a faculty job is a very isolating process, only few postdocs that I personally knew were sailing in the same boat. | (1) |
| **Frustrating** | It was a little frustrating to no receive rejection letters from the majority of institutions.<br>- I was really frustrated when I didn't hear from a place after I did an in-person interview for 2 &1/2 months | (1) |
| **Unpredictable** | I found it very unpredictable whether the institutes were interested in my profile or not. Also the application procedures varied a lot, in terms of the number of rounds, time frame and information updates. | (1) |
| **Information-Sparse** | Most places do not indicate how many pages they want each of the documents they are requesting. | (2) |
| **Non-ordinary requests** | Some schools made it particularly difficult by having non-ordinary requests, such as personal statements, 5 reference letters (instead of 3), 3 separate research proposals etc. | (1) |

| | | |
|---|---|---|
| **Black-box** | The community of other postdocs on the market helping each other deal with questions and anxieties really helped to demystify the black box that is the faculty search. | (1) |
| **Burden-on-research** | The application process was extremely time-consuming during my postdoctoral research. | (1) |

**Table S22.** Overview of candidates who commented on their view in general of the application process. Responses to "Do you have any comments that you would like to share? For example, how did you experience the application process?" Survey participants were able to provide long answers to this comment question. A word cloud referring to this table of comments is provided in Figure 7C.

| **Table S23. Applicant responses to "Why did you find the Future PI Google Sheet helpful?** | |
|---|---|
| **Theme (frequency of responses)** | **Survey Responses** |
| **Very helpful or helpful (30)** | The Future PI slack channel/spreadsheet was really helpful to guide me during the process providing real time feedback of what's going on in the job market in comparison to advice from PIs that experienced years ago (or that experienced in a completely different way that I did). |
| | It helped me realize my status with more than 70% of the universities that I had applied to. This is a much better approach (quick painless death to a school I have not been called for, but others have) than patiently waiting, hoping against hope as the months drag by. |
| | Did not find the Future PI google sheet particularly helpful because there is almost no one else on there in my field. But I did find the Future PI slack discussions helpful. The future PI google sheet was super helpful to ask questions, vent, cross out schools that ghosted me but others have heard back from, and just overall learn a ton of tips on navigating this process. |
| | Found it useful to keep track of whether other people had heard back from schools I applied to or not. |
| | Used Future PI google sheet helpful but also stressful. I also think it's heavily biased towards those who are successful on job market- thus not a good picture of the actual process. I think >90% of people who filled out the application stats had at least one onsite visit. I do not think this is indicative of most people success rate. |
| | The Future PI Slack and spreadsheet was very helpful for tracking interviews and asking advice. |
| | Checked the future PI spreadsheet almost every day. It was almost like a soulmate during the ridiculously stressful process. I never realized that so |

| | |
|---|---|
| | many people have similar anxieties, insecurities, and questions as me. Applying for a faculty job is a very isolating process, only few postdocs that I personally knew were sailing in the same boat. Access to the information & comments posted there is precious. It was a true collaborative effort by many anon. contributors, I hope it stays that way, and no one takes control or credit for the resources posted there. |
| | Future PI google sheet and slack group were both extremely helpful. |
| | Really useful because at least you know if someone got selected for the position. |
| | The Future PI google sheet was a great resource for understanding the timing of things, how many applications people submitted, and general advice/feedback. |
| | It was helpful to know if others had heard from places I ad not as well as get a sense of the general competitiveness of the market. |
| | Found the Future PI google sheet very helpful to know when other people heard back from jobs I had applied to. |
| | Found the google sheet very helpful, though not many of the listings were in my field it was nice to see the expected timings from one step to the next, and see where my search landed in terms of response rates. |
| | The Future PI google sheet and it was incredibly helpful. The community of other postdocs on the market helping each other deal with questions and anxieties really helped to demystify the black box that is the faculty search. |
| | Future PI google sheet was extremely useful to track the status of jobs and to ask for advice. |
| | It was great for moral support. |
| | It was extremely helpful to not feel so alone in the process. |

**Table S23.** Overview of candidates who were familiar with the Future PI Slack resource and other resources during their application process. Responses to “Why did you find the Future PI google sheet/Slack helpful?” Survey participants were able to provide long answer to this comment question (Future PI Slack of FPI Slack is a slack group comprised of postdoctoral researchers aspiring to apply for faculty positions).

| Table S24. Twitter Poll #2: Researcher Time Spent on faculty job applications | | | | |
|---|---|---|---|---|
| **Theme** | **Less than 1 hour** | **1-2 hours** | **2-3 hours** | **Greater than 3 hours** |
| **Number of Respondents (n=234)** | 6% (n=14) | 15% (n=35) | 16% (n=37) | 63% (n=147) |

**Table S24.** Overview of the responses to a Twitter poll with the question: "How long on average did you, the applicant, spend preparing each faculty job application?".

| Table S25. Twitter Poll #3: Effect of time spent on faculty applications on research | | | |
|---|---|---|---|
| **Theme** | **Yes** | **No** | **Just show me the poll results** |
| **Number of Respondents (n=164)** | 58% (n=95) | 8% (n=13) | 34% (n=56) |

**Table S25.** Overview of the responses to a Twitter poll with the question: "Do you feel like time spent preparing your faculty job applications impaired your ability to push other aspects of your career forward? (pubs, grant apps, research goals)".

| Table S26. Themes from Job Applicant survey written responses to helped your application | | |
|---|---|---|
| **Theme** | **Survey Responses** | **Frequency of the response *n*** |
| **Networking** | Built an extensive network of colleagues and collaborators: future colleagues lobbied heavily for me | (4) |
| | Attending many meetings, hugely important in my success | (4) |
| | Personal connections seemed helpful and fit seemed really important | (4) |
| **Preprints** | Preprints were enormously helpful: interview based on pre-print, offer based on preprint, helpful to show adopting new techniques and productivity, was looked favorably upon much before paper published | (4) |
| **Publications** | Paper being accepted was important | (5) |
| | High-impact non-CNS publications | (1) |

| | | |
|---|---|---|
| | CNS publications helped get my position | (2) |
| **Research Field** | Performing interdisciplinary research | (1) |
| | Learning a new research technique | (1) |
| | Research area being perfect fit for department: asking hiring departments what/which field/expertise they are looking for | (4) |
| **Mentoring** | Mentoring goals align well with those of the university | (3) |
| | Having mentors helps you get through this process | (1) |
| | Having colleagues/mentors (both early career and senior) review and comment on application | (3) |
| | Significant amount of leadership experience: through helping to start an internship for underrepresented minority undergraduates | (1) |
| **Teaching** | Teaching experience being perfect fit for department | (1) |
| | Teaching certificate was immensely helpful for teaching focused positions. | (1) |
| **Funding** | Having funding in the past: e.g. Postdoctoral Fellowship (NIH F32 fellowship) and K awards, Marie Curie fellowship, Human Frontier fellowship were looked favorably upon | (12) |
| | Institutional Research and Academic Career Development Award | (1) |
| | Experience of writing grants: co-written with PI, scored not funded | (1) |
| **Pedigree** | PhD and/or Postdoctoral lab pedigree | (3) |
| **Service** | Serving as a reviewer for journals | (1) |
| **Other Skills** | Other skills are also important not just research skills | (1) |
| **Randomness** | The process of applying and finding a job seems completely up to luck/lottery, that there is no magic formula to this, **but having amazing mentors helps you get through this process**. | (3) |

**Table S26.** Candidate responses to “Was any aspect of your career particularly helpful when applying (preprints, grants etc.)?” Survey participants were able to provide long answers to this comment question. A word cloud referring to this table of comments is provided in Figure 7A.

| Table S27. Themes from written Applicant responses to obstacle for their application | | |
|---|---|---|
| **Theme** | **Survey Responses** | **Frequency of the response *n*** |
| **No application feedback** | Hard to say because I got no feedback, More transparency from the search committee would have been great. | (5) |
| **Nepotism** | Nepotism based on advisor name | (3) |
| | Nepotism based on institution pedigree | (1) |
| | Same finalists invited everywhere: all the top schools only invited these anointed candidates for site interviews, Lack of breadth and diversity in the pool of on-site candidates | (1) |
| **Personal Life concerns** | Maternity leave | (3) |
| | Two-body problem (partners finding jobs in the same vicinity) | (1) |
| | Work-Family balance (balancing care for children with time-consuming and stressful faculty job applications) | (2) |
| **Poor Mentorship** | No mentor that gave feedback on writing grants | (8) |
| | Lack of mentorship on how to handle the on-site interview and structure my proposed work talk. | (1) |
| **Publications** | Lack of postdoc papers being accepted | (3) |
| | lack of CNS papers | (4) |
| | harmful was not having postdoc papers all in preparation and not published yet | (1) |
| | publication gap following the birth of my first child was a problem | (2) |
| | Maintaining productivity in the lab while preparing applications was difficult. | (1) |
| **Preprints** | Lack of Preprints | (1) |
| **Citizenship** | Not being a US citizen made it hard to apply for the majority of fellowships or the NIH K or F type awards | (3) |

| | | |
|---|---|---|
| **Language Skills** | Level of speaking English is a big hurdle. | (1) |
| **Research Field** | No offers due to *in vitro* work instead on *in vivo* research (lack of fit) | (1) |
| | Interdisciplinary research was underappreciated | (6) |
| | Not Moving Institutions from PhD to postdoc | (1) |
| | Working in industry for a few years before applying to academic position. | (1) |
| **Teaching** | Lack of teaching experience that would have also helped with writing teaching statements | (2) |
| **Funding** | Lack of Funding: No funding, ability to procure outside funding | (8) |
| **Randomness/Chaos** | The academic market is 90% chaos and privilege. It's like the lottery, but worse. | (1) |

**Table S27.** Candidate responses to “Was any aspect of your career particularly an obstacle when applying?” Survey participants were able to provide long answers to this comment question. A word cloud referring to this table of comments is provided in Figure 7B.

| **Table S28. Frequency of Job Applicant comments who received an offer** | | |
|---|---|---|
| **Type of Candidates** | **Percentage of Candidates** | **Nature of the Comments** |
| All Candidates | 100%<br>(317 out of 317) | - |
| Candidates with at least one offer | 58.3%<br>(185 out of 317) | - |
| Candidates with at least one offer who wrote a comment about the application process | 9.2%<br>(17 out of 185) | - |
| Candidates with offers who wrote a comment and had **positive** perception of the application process | Less than 1.8%<br>(2 out of 17) | e.g. Fine, Smooth |
| Candidates with offers who who wrote a comment and had **negative** perception of the application process | 88.2%<br>(15 out of 17) | e.g. Terrible, Stressful |
| Candidates with offers who had **positive** perception of the application process | 1.1%<br>(2 out of 185) | - |

| | | |
|---|---|---|
| Candidates <u>with offers</u> who had **negative** perception of the application process | 8.1%<br>(15 out of 185) | - |
| Candidates <u>with offers</u> who made **no comments** | 61.1%<br>(113 out of 185) | - |
| Candidates <u>with or without offers</u> who made **no comments** | 63.4%<br>(201 out of 317) | - |

**Table S28.** Overview of candidates who commented on their view in general of the application process. Responses to "Do you have any comments that you would like to share? For example, how did you experience the application process?" Survey participants were able to provide long answer to this comment question. A word cloud referring to this table of comments is provided in Figure 7C. Percentages are calculated out of the total number of respondents to this particular survey questions.

**Table S29. themes from Search committee written responses to any Other comments or thoughts about the state of hiring for tenure track positions?**

| Theme | Survey Responses | Frequency of the response *n* |
|---|---|---|
| **Extending a faculty job offer is not trivial.** | If you took the probability of an offer an applied it blindly to the number of applicants, you would conclude that it is nearly hopeless to apply. The reality is that a small number of folks end up with multiple offers. So it is either more or less hopeless, depending on whether you're one of that small number. In my experience, it is easy to imagine that it's all luck or whose lab you come from or whether you got a CNS paper, this attitude evaporates when you experience an actual faculty hiring process. Applicants' track records are nontrivially long, and they are subjected to many forms of poking and prodding to see if they're for real before an offer is extended. | (1) |
| **Quality of publications are most important.** | The quality of the candidate's published research is the most important thing we try to evaluate. | (1) |
| **Candidate perceptions of the hiring process are unreal.** | The perception gap between what trainees think matter and what actually matters is rather large due to a number of correlations. There are many problems with academic hiring processes, but | (1) |

| | | |
|---|---|---|
| | they are typically not the ones trainees think they are. | |
| **Too many candidates do not aim for a fit with their ability.** | Too many candidates don't aim correctly for where they actually fit with their record and ability - aiming either too high or too low. We have limited slots for interviews, and often don't bring in our very top applicants since they invariably turn us down for a top 5 place. | (1) |
| **Too many candidates apply for faculty positions.** | There are too many people applying for tenure track jobs. Postdocs should think more carefully about what it is they truly like about research and look for jobs that allow them to do that. | (1) |
| **Finding a fit between candidate and the department is key.** | Most find a home. Process is about connecting with a community that makes sense. | (1) |
| **Candidates apply to too many positions** | Social media has amplified the crazy nature of the process. People apply to too many positions. | (1) |
| **Finding a faculty job requires persistence** | It is not easy and requires persistence. | (1) |
| **Challenges in Finding a Fit** | Although we have many fantastic applicants each cycle, it remains challenging to and someone who is an excellent intellectual fit for our department AND has superlative credentials. | (1) |
| **The hiring process is not as bleak as portrayed.** | It's not as bleak as the scientific press or many postdocs think. | (1) |

**Table S29.** Overview of search committee members who commented on "Do you have any other comments or thoughts about the state of hiring for tenure track positions?" Survey participants were able to provide long answer to this comment question.

| **Table S30. Applicant Demographics: Number of times (cycles/years) that the candidates had applied for faculty positions** | |
|---|---|
| **Theme** | **Applicant number** |
| **Total Number of Applicants** | 100% (317 out of 317) |
| **Applied 0 cycles** | Less than 1% (1 out of 314) |

| | |
|---|---|
| **Applied 1 cycle only** | 56.7% (176 out of 314) |
| **Applied for 2 cycles so far** | 29.3% (92 out of 314) |
| **Applied for 3 cycles so far** | 10.5% (33 out of 314) |
| **Applied for 4 cycles so far** | 3.2% (10 out of 314) |
| **Applied for 5 cycles** | Less than 1% (2 out of 314) |
| **Did not respond to this survey Question** | Less than 1% (3 out of 317) |

**Table S30.** Overview of number of times job candidate survey respondents applied for a faculty (PI) position (Box 1). This is in response to the survey question :"How many times have you applied for PI positions? i.e. if the 2018-2019 cycle was the first time, please enter "1", if you also applied last cycle, enter "2", etc. Percentages are calculated out of the total number of respondents to this particular survey questions (n=314).

| **Table S31. Statistics from the Search Committee Survey** | |
|---|---|
| **Theme** | Faculty Responses (n=15) |
| **Typical number of applications committees received each cycle** | 100-199 applications (n=5)<br>200+applications (n=10) |
| **Typical number of applicants making through the first round of cuts** | 1-19 candidates (n=5)<br>20-39 candidates (n=4)<br>40-50 candidates (n=3)<br>60+ candidates (n=3) |
| **Typical number of applicants invited for offsite interview (Skype/phone)** | 0 candidates (n=4)<br>5 or fewer candidates (n=1)<br>6-8 candidates (n=1)<br>5-10 candidates (n=1)<br>8-10 candidates (n=1)<br>10 candidates (n=2)<br>10-15 candidates (n=1)<br>12-15 candidates (n=2)<br>16 candidates (n=1)<br>NA (n=1) |
| **Typical number of applicants invited for onsite (on campus) interview** | 5 candidates (n=5)<br>6 candidates (n=4)<br>8 candidates (n=5)<br>10 candidates (n=1) |
| **Typical number of offers committees make per job posting** | 0-1 candidates (n=11) |

| | |
|---|---|
| | 2-3 candidates (n=4) |
| **Number of openings at your department in the last five years** | 2-3 openings (n=5)<br>4-5 openings (n=6)<br>6+ openings (n=4) |
| **Length of faculty involvement with department's search committee** | 1- 4 years (n=1)<br>5-10 years (n=3)<br>11-19 years (n=8)<br>20-29 years (n=2)<br>30+ years (n=1) |

**Table S31.** Overview of the search committee survey responses to "Approximately how many applicants for a posted position do you get?", "Approximately how many applicants make it through the first round of cuts?", "Approximately how many applicants are invited for off-site interview (Skype/phone)?", "Approximately how many offers does your committee make per job posting?", "Approximately how many openings has your department had in the last five years?", "Approximately how many applicants are invited for on-site interview?", "How long have you been involved in academic search committees?".

| **Table S32. Search Committee survey demographics** | |
|---|---|
| **Theme: Faculty field and institution type** | **Faculty number** |
| **Total Number of Faculty responses received** | **(All faculty were based in the United States n=15)** |
| **Number of Faculty involved with search committees in the past 10 years** | 60% (9 out of 15) |
| **Faculty in Engineering** | 6.7% (1 out of 15) |
| **Faculty in Life Sciences** | 87% (13 out of 15) |
| **Faculty in Chemistry** | 6.7% (1 out of 15) |
| **Work at R1: Doctoral Universities** | 100% (15 out of 15) |

**Table S32.** Overview of the search committee faculty demographics of our faculty survey respondents. Percentages are calculated out of the total number of respondents to this particular survey questions (n=15).

| Table S33: Statistics from the Search Committee Survey on Preprints | | | | | |
|---|---|---|---|---|---|
| **Theme** | **Depends on the faculty member. Some love preprints, others do not.** | **Yes (preprints are appreciated and considered a demonstration of productivity)** | **No (preprints are largely ignored)** | **We have not searched in the last 2 years** | **Appreciated and used but not as heavily weighted as peer-reviewed published work.** |
| **Does your committee look favorably upon preprints? (n=15)** | 6.6% (1 out of 15) | 66.7% (10 out of 15) | 13.3% (2 out of 15) | 6.6% (1 out of 15) | 6.6% (1 out of 15) |

**Table S33.** Overview of the search committee survey responses to "Does your committee look favorably upon preprints?".

| Table S34. Statistics from the Search Committee Survey on Perception of the job market | |
|---|---|
| **Theme** | **Faculty perception of the job market for tenure-track faculty as someone involved in the search process (please tick all that are true) (n=15)** |
| **Candidates fall below expectations during interviews** | 66.7% (10 out of 15) |
| **Too few good applicants** | 13.3% (2 out of 15) |
| **Hard to identify good candidates from applications** | 13.3% (2 out of 15) |
| **Easy to identify good candidates from applications** | 73.3% (11 out of 15) |
| **Too many good applicants** | 66.7% (10 out of 15) |
| **Candidates surpass expectations during interviews** | 626.6% (4 out of 15) |
| **Skype interviews have helped reduce mismatch** | 6.6% (1 out of 15) |
| **The Market has changed alot since I applied for a faculty position** | 20% (3 out of 15) |

**Table S34.** Overview of the search committee survey responses to "What is your perception of the job market for tenure track faculty as someone involved in the search process (please tick all that are true)". Percentages are calculated out of the total number of respondents to this particular survey questions (15).

| Table S35. Statistics from the Search Committee Survey on weighting of applicant materials | |
|---|---|
| **Theme** | **Faculty evaluation (on a scale of 1-5) of the various tenure track application materials (total committee members n=15)** |
| **Weight of prior teaching experience** | 0% (0 9 out of 15) said 5<br>0% (0 out of 15) said 4<br>20% (3 out of 15) said 3<br>53.3% (8 out of 15) said 2<br>26.7% (4 out of 15) said 1 |
| **Weight of good mentorship in the candidate's postdoctoral/graduate student lab explicitly on selection process (e.g. "This candidate's mentor is known to produce good trainees")** | 6.7% (1 out of 15) said 5<br>20% (3 out of 15) said 4<br>26.7% (4 out of 15) said 3<br>40% (6 out of 15) said 2<br>6.7% (1 out of 15) said 1 |
| **Weight of the research proposal weigh on the selection process (e.g. "This candidate's research statement is incredibly compelling!")** | 60% (9 out of 15) said 5<br>33.3% (5 out of 15) said 4<br>0% (0 out of 15) said 3<br>6.7% (1 out of 15) said 2<br>0% (0 out of 15) said 1 |
| **Weight of the graduate student fellowships or awards (e.g. NSF GRF, NIH F30, etc.)** | 6.7% (1 out of 15) said 5<br>20% (3 out of 15) said 4<br>33.3% (5 out of 15) said 3<br>33.3% (5 out of 15) said 2<br>6.7% (1 out of 15) said 1 |
| **Weight of the non-transitional postdoctoral fellowships or awards (e.g. NIH F32, AHA etc.)** | 20% (3 out of 15) said 5<br>33.3% (5 out of 15) said 4<br>33.3% (5 out of 15) said 3<br>13.3% (2 out of 15) said 2<br>0% (0 out of 15) said 1 |
| **Weight of the transition awards as a positive factor (i.e. K99/R00 award, Burroughs Wellcome Career Award, or another award that provides the applicant with money as a new faculty member)** | 33.3% (5 out of 15) said 5<br>20% (3 out of 15) said 4<br>33.3% (5 out of 15) said 3<br>13.3% (2 out of 15) said 2<br>0% (0 out of 15) said 1 |
| **Weight of the Cell, Science, or Nature papers above papers in other journals** | 0% (0 out of 15) said 5<br>26.7% (4 out of 15) said 4 |

| | |
|---|---|
| | 13.3% (2 out of 15) said 3<br>26.7% (4 out of 15) said 2<br>33.3% (5 out of 15) said 1 |
| **Weight of the journal impact factor explicitly in to the selection process (e.g. does the word 'impact factor' come up in discussions around applicants)** | 0% (0 out of 15) said 5<br>13.3% (2 out of 15) said 4<br>6.7% (1 out of 15) said 3<br>26.7% (4 out of 15) said 2<br>53.3% (8 out of 15) said 1 |

**Table S35.** Overview of the search committee survey responses to evaluation of a number of the tenure-track application materials: 1) "To what extent does the research proposal weigh on the selection process (e.g. "This candidate's research statement is incredibly compelling!", 2) "To what extent does good mentorship in the candidate's postdoctoral/graduate student lab explicitly weigh on selection process (e.g. "This candidate's mentor is known to produce good trainees", 3) "How heavily does the committee weigh graduate student fellowships or awards (e.g. NSF GRF, NIH F30, etc.)", 4) "How heavily does the committee weigh non-transitional postdoctoral fellowships or awards (e.g. NIH F32, AHA etc.)", 5) "Does your committee weigh Cell, Science, or Nature papers above papers in other journals?", 6) "To what extent does journal impact factor explicitly weigh in to the selection process (e.g. does the word 'impact factor' come up in discussions around applicants)?", 7) "How heavily does the committee weigh transition awards as a positive factor (i.e. K99/R00 award, Burroughs Wellcome Career Award, or another award that provides the applicant with money as a new faculty member)?", 8) "How heavily does the committee weigh prior teaching experience?". In the survey, a 5-level Likert scale was used to record faculty impressions where a response of 1=not at all and 5=heavily. Percentages are calculated out of the total number of respondents to this particular survey questions (n=15).

**Table S36. Themes from the Search Committee Survey written responses to: What information do you wish more candidates knew when they submit their application?**

| Theme | Survey Responses | Frequency of the response |
|---|---|---|
| **Importance of Research Proposal** | In a useful sense, no one cares about your particular interests in the way that you do. Write your application to engage a broad group of scientists. We care about the impact of what you propose to do (research blurb). | (5) |
| **Importance of publications** | That your research and your papers are the main thing we look at, and we actually do read and evaluate your work. Where it's published (bioRxiv vs. CNS) does not matter. We're looking for leading indicators of success, not lagging ones. | (2) |

| | | |
|---|---|---|
| **Importance of the Chalk-Talk** | We care about whether you can tell us how and why this matters (chalk talk). | (3) |
| **Importance of the Impact of your work** | We care about the impact of what you have done | (2) |
| **Importance of being a great & creative colleague** | I wish candidates better understood that we are looking for an interesting colleague. | (3) |
| **Importance of being independent** | Evidence for independence innovative and creative research plans are good. | (1) |
| **Importance of being authentic** | Making authentic personal connections when interviewing is always crucial. | (1) |

**Table S36.** Overview of Search Committee who responded to "What information do you wish more candidates knew when they submit their application?" Survey participants were able to provide long answer to this comment question. A word cloud referring to this table of comments is provided in Figure 9a.

| **Table S37. Themes from Search Committee write in responses to any changes in the search process since the first search you were involved in?** | | |
|---|---|---|
| **Theme** | **Survey Responses** | **Frequency of the response *n*** |
| **No Change** | Not much/Nothing fundamental | (3) |
| **Has changed** | Yes | (2) |
| **Negative publicity** | There is a lot more negative publicity and discussion. | (1) |
| **Higher credentials required** | The credentials required to pass the first cut are getting higher and higher. | (1) |
| **Stronger Candidates** | The candidates are stronger/more savvy now | (2) |
| **Addition of online interviews** | The Skype/Zoom off-site interview component is new compared to six years ago when I interviewed. | (3) |
| **Recommendation Letters** | Letters have become more hyperbolic | (1) |
| **Chalk-Talks** | The chalk talk is a bigger deal now | (2) |

**Table S37.** Overview of search committee faculty members who commented on “Have you noticed any changes in the search process since the first search you were involved in?” Survey participants were able to provide long answer to this question. A word cloud referring to this table of comments is provided in Figure 9.

| **Table S38. Logistic Regression analysis on the job applicant survey data** | | | | | | | | |
|---|---|---|---|---|---|---|---|---|
| **VARIABLE** | **Coefficient** | | **P-Value** | | **STD. Error** | | **Z-Value** | |
| | **Missing data excluded** | **Missing data imputed** | **Missing data excluded** | **Missing data imputed** | **Missing data excluded** | **Missing data imputed** | **Missing data excluded** | **Missing values imputed** |
| **(Intercept)** | 0.2650 | **0.5158** | 0.296 | $1.12 \times 10^{-4}$ | 0.2534 | 0.1336 | 1.0455 | 3.8623 |
| **Female applicants** | 0.4055 | **0.3039** | 0.118 | $3.29 \times 10^{-2}$ | 0.2596 | 0.1424 | 1.5622 | 2.1337 |
| **Field of research** | -0.0597 | -0.2120 | 0.773 | 0.113 | 0.2074 | 0.1337 | -0.2878 | -1.5858 |
| **CNS (first authorship)** | 0.1963 | 0.1995 | 0.473 | 0.165 | 0.2734 | 0.1437 | 0.7178 | 1.3884 |
| **CNS (co-authors hip)** | -0.6758 | 0.1939 | 0.161 | 0.172 | 0.4824 | 0.1420 | -1.4010 | 1.3658 |
| **Application number** | **0.4874** | **0.6411** | $4.42 \times 10^{-2}$ | $1.23 \times 10^{-4}$ | 0.2422 | 0.1669 | 2.0122 | 3.8402 |
| **Other jobs** | -0.2362 | **-0.3477** | 0.320 | $9.76 \times 10^{-3}$ | 0.2373 | 0.1345 | -0.9955 | -2.5843 |
| **Funding** | -0.1953 | **0.3913** | 0.406 | $5.86 \times 10^{-3}$ | 0.2351 | 0.1420 | -0.8308 | 2.7555 |
| **Patent** | -0.1704 | -0.0154 | 0.578 | 0.914 | 0.3063 | 0.1424 | -0.5563 | -0.1083 |

| | | | | | | | | |
|---|---|---|---|---|---|---|---|---|
| **Total number of publications** | 0.1441 | -0.4599 | 0.777 | 0.119 | 0.5082 | 0.2948 | 0.2837 | -1.5600 |
| **Number of first-author publications** | -0.0547 | 0.1786 | 0.870 | 0.376 | 0.3337 | 0.2018 | -0.1639 | 0.8854 |
| **Postdoc fellowships** | 0.3945 | 0.2244 | $9.24 \times 10^{-2}$ | $9.33 \times 10^{-2}$ | 0.2344 | 0.1337 | 1.6829 | 1.6783 |
| **PhD fellowships** | -0.3604 | -0.1506 | 0.156 | 0.272 | 0.2542 | 0.1370 | -1.4180 | -1.0991 |
| **Adjunct teaching experience** | 0.3708 | 0.3126 | 0.162 | $7.86 \times 10^{-2}$ | 0.2654 | 0.1777 | 1.3968 | 1.7586 |
| **Citations** | 0.3811 | 0.1252 | 0.315 | 0.611 | 0.3796 | 0.2461 | 1.0040 | 0.5086 |
| **h-index** | 0.2878 | **0.6709** | 0.500 | $1.85 \times 10^{-2}$ | 0.4268 | 0.2848 | 0.6744 | 2.3556 |
| **Years on job market** | -0.1657 | **-0.2887** | 0.436 | $3.68 \times 10^{-2}$ | 0.2126 | 0.1383 | -0.7796 | -2.0879 |

**Table S38.** Regression analysis was performed on the job applicant survey data. All variables collected except for number of remote and on-site interviews were included as potential predictors of receiving (1) or not receiving (0) a job offer. Positive coefficients indicate positive associations and negative coefficients indicate negative associations with receiving an offer. Coefficients that are zero indicate no association. Bold values indicate that the associations were found to be significant at a threshold of 0.05

| Table S39. Job Applicant survey scholarly metrics by gender with breakdown by offer status | | | | | | | | | | | | | | | |
|---|---|---|---|---|---|---|---|---|---|---|---|---|---|---|---|
| **Variable** | **All (Fig. 2B)** | | | | | **With offers** | | | | | **Without offers** | | | | |
| | **Mean** | | **Median** | | **p-value** | **Mean** | | **Median** | | **p-value** | **Mean** | | **Median** | | **p-value** |
| | **F** | **M** | **F** | **M** | | **F** | **M** | **F** | **M** | | **F** | **M** | **F** | **M** | |
| **Number of first-author publications** | 6.1 | 7.8 | 5 | 7 | **$1.42 \times 10^{-4}$** | 6.0 | 8.6 | 5 | 7 | **$1.92 \times 10^{-5}$** | 6.4 | 6.8 | 5 | 7 | 0.409 |
| **Total number of publications** | 13.5 | 16.4 | 11 | 14 | **$2.79 \times 10^{-3}$** | 13.5 | 17.5 | 11 | 16 | **$2.68 \times 10^{-4}$** | 13.4 | 14.9 | 11 | 12 | 0.570 |
| **Citations** | 397.0 | 455.3 | 228 | 343 | **$1.50 \times 10^{-2}$** | 453.0 | 522.0 | 251 | 403 | **$3.18 \times 10^{-2}$** | 315.4 | 358.4 | 216.5 | 265 | 0.233 |
| **h-index** | 7.9 | 9.0 | 7 | 9 | **$5.42 \times 10^{-3}$** | 8.1 | 9.9 | 7 | 10 | **$9.42 \times 10^{-4}$** | 7.7 | 7.8 | 7 | 8 | 0.777 |

**Table S39.** Mean and median values for publication-related metrics plotted in Figure. 2B broken down by gender and offer status. Additionally, p-values from Wilcoxon rank-sum tests that compare metric values from the female and male groups. “All” shows these values when the full dataset is considered, “With offers” shows values for only those applicants with at least one offer, and “Without offers” shows values for only those without any offers. “F” stands for female and “M” stands for male. Trends in gender differences remain the same even for the applicants with offers, serving as a possible explanation for the similar search outcomes for females and males and the importance of gender in the logistic regression.

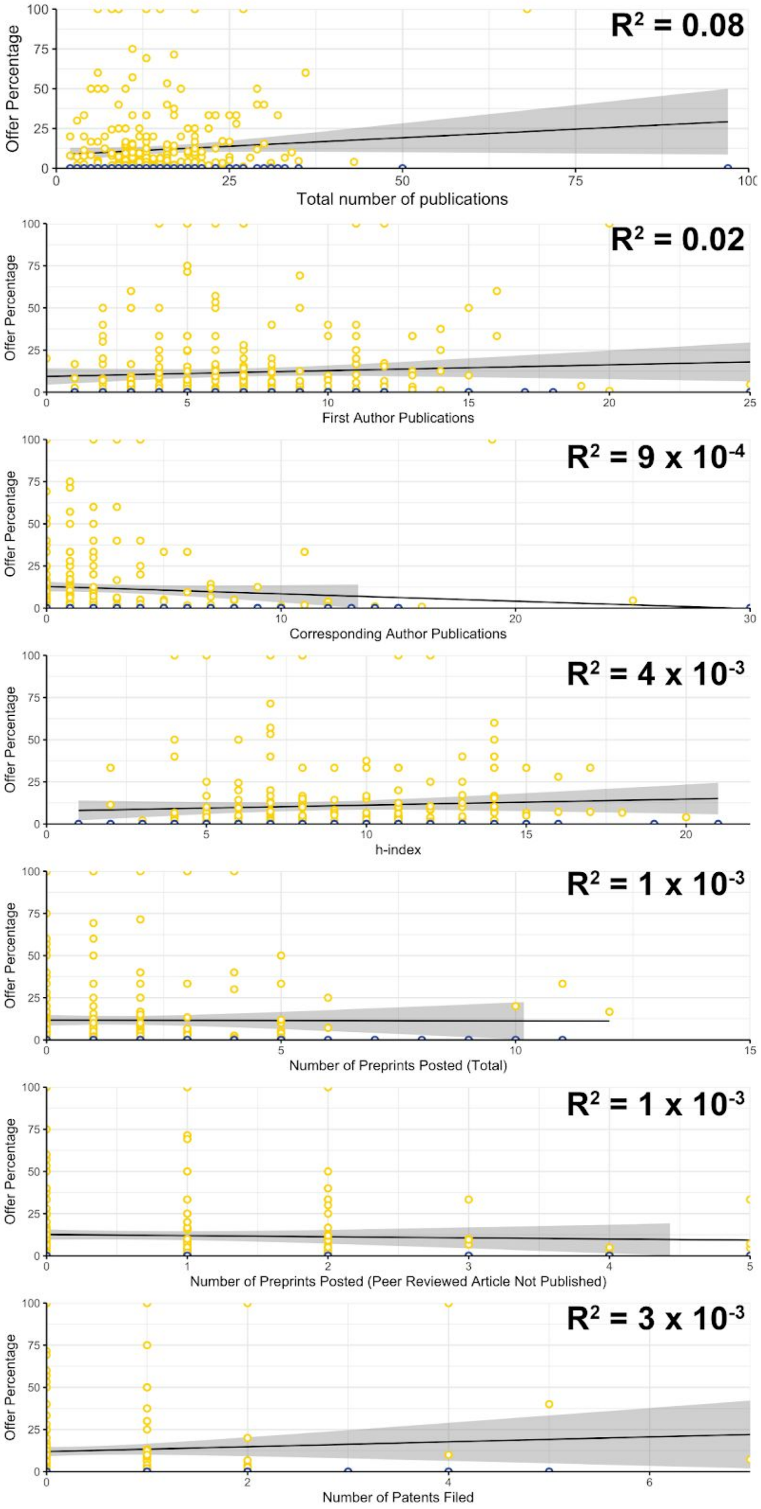

**Figure S1.** Correlation ($R^2$ = Pearson correlation coefficient) between offer percentage and a number of traditional metrics: total number of publications (top), number of first author publications (2nd graph), number of corresponding author publications (3rd graph), h-index (4th graph), preprints posted (overall total, 5th graph; as well as those in which the peer-reviewed article was not published at the time of application, 6th graph), and number of patents filed, bottom graph). Yellow dots represent candidates with an offer, blue dots received no offers; black line represents linear best-fit and gray fill represents the 95% confidence interval for that fit.

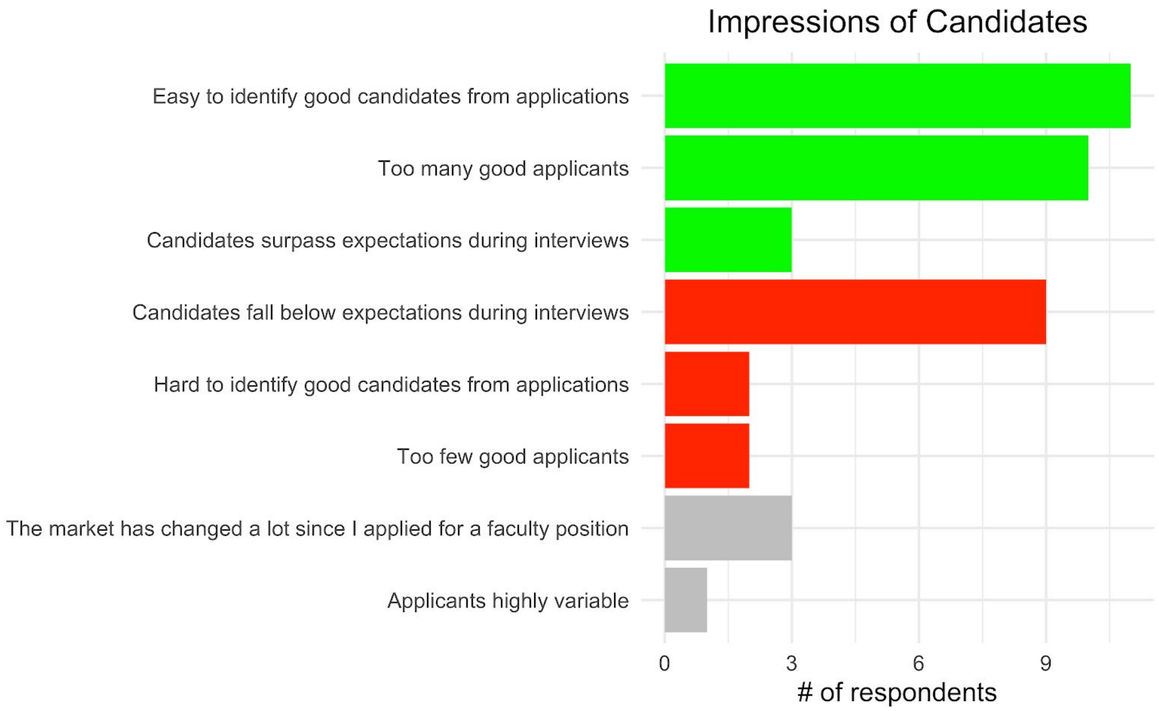

**Figure S2.** Overview of the search committee impressions of the candidates (refers to Table S34).

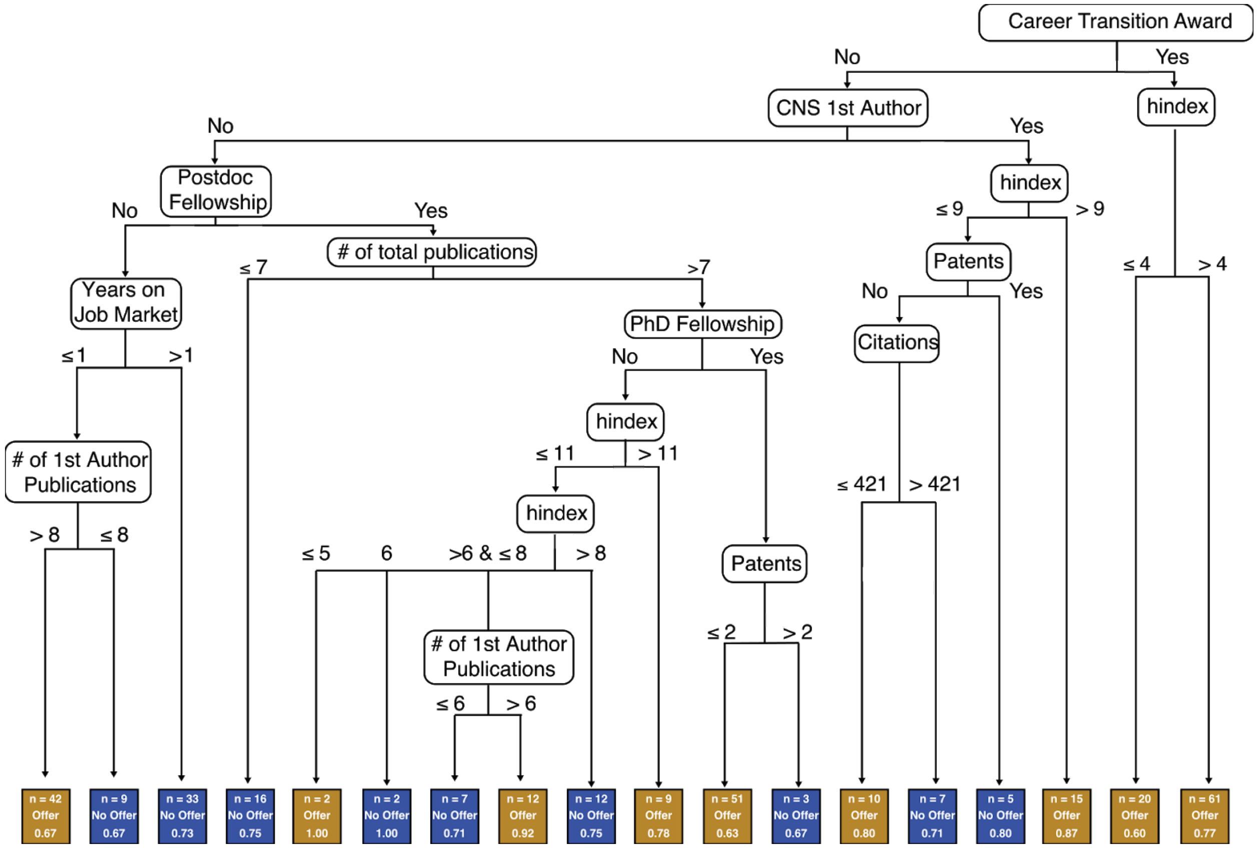

**Figure S3.** Visualization of possible paths to an offer using the C5.0 decision tree algorithm. Each rounded node represents an independent variable and each rectangular note represents one of two possible outcomes (offer (gold) or no offer (blue)). Only those variables in Figure 4B were included. In the case of binary variables such as funding and fellowships, “>0” indicates a “yes” and “<=0” indicates a “no”. All other variables, except for h-index, were split based on counts. The outcome nodes are labeled with three pieces of information: (1) the number of applicants who fell into the given branch (*n*), (2) the most common outcome in that branch, and (3) the fraction of individuals with that outcome. For example, the rightmost branch shows applicants who had a career transition award and h-index > 4. They constitute the largest group in our dataset (61 individuals). However, only 77% of these applicants received an offer. Similarly, the second and third largest groups included 51 applicants (63% with offer) and 42 applicants (67% with offer) respectively (see eighth outcome box from right and leftmost box). These three groups accounted for 48.6% of our survey respondents. Note that while decision trees have often been used as prediction models, this tree is only reflective of our dataset and choice of algorithm and parameters. We have used this solely for visualization purposes and advise against using this prospectively to evaluate chances of success on the job market as there may be alternative trees that are equally plausible and accurate. In fact, the accuracy of the overall decision tree in distinguishing between candidates with offers and those without was only 58.5%. Furthermore, no group with more than two applicants consisted purely of those with offers and those without. Even in the nine groups where the most common outcome was “no offer”, on average, 25% of the applicants did receive offers.

#### The job Applicant survey

##### Survey of the applicants to the tenure-track jobs

Please answer in as much detail as you can. All responses will be treated fully anonymously.
*Required

1. Where (which country) did you apply for faculty positions?*
   United States
   Canada
   United Kingdom
   Germany
   Switzerland
   Australia
   Singapore
   Other: Short Answer---------------

2. Your field (broadly defined)*
   Biomedical or life sciences
   Bioengineering
   Biology (other)
   Chemistry
   Chemical engineering
   Computer Science
   Mechanical engineering
   Physics
   Other: Short answer----------------

3. Did you apply primarily to research (R1) or teaching-focused (PU) institutions?
   R1 Institutions
   PUIs
   Both R1 & PUIs
   Other: Short answer----------------

4. Total Number of applications submitted June 2018-April 2019*
   Short Answer:---------------------

5. Number of remote/offsite (e.g. phone, Skype...) interviews*
   Short Answer:---------------------

6. Number of onsite interviews*
   Short Answer:---------------------

7. Number of offers*
   Short Answer:---------------------

8. Approximate Number of rejections
   Short Answer:---------------------

9. Do you have any comments that you'd like to share? For example, how did you experience the application process? Did you use the Future PI google sheet? Did you find it helpful? If yes, why?
   Long Answer:---------------------

10. Where are you currently working (country)?
    Short Answer---------------

11. What's your gender?
    Male
    Female
    Non-Binary
    Prefer not to disclose
    Other: Short Answer---------------

12. What is your current position?
    Postdoc (or equivalent, if it is a postdoc position with a different title)
    PhD student
    Other: Long Answer-----------------

13. If postdoc, how many years have you been a postdoc? (TOTAL number of years, if you've had multiple appointments.)
    Short answer:----------------------

14. If postdoc, is this your first postdoc position?
    1st postdoc
    2nd postdoc
    >2nd postdoc

15. How many times have you applied for PI positions? I.e. if the 2018-2019 application cycle was the first time, please enter "1", if you also applied last cycle, enter "2", etc.
    Short Answer:---------------------

16. If you have a Google Scholar account, what is your number of citations (all; since 2014), and h-index (all; since 2014)
    Short Answer:---------------------

17. How many papers have you published? (ALL papers: as co-author, first author and last author. If conference abstracts count towards publications in your field, please include those as well.)
    Short Answer:---------------------

18. How many first author papers have you published?
    Short Answer:---------------------

19. How many papers have you published as corresponding author?
    Short Answer:---------------------

20. What is your highest impact-factor publication (IF or journal)? Was this as first author, a co-author, or as corresponding author? (Suggested format for answer: journal name, author description) [e.g. eLife, first author]
    Short Answer:---------------------

21. Did you have any Cell/Nature/Science publications? If yes, how many and were you first author, a co-author, or corresponding author on them? (Suggested format for answer: number, author description) [e.g. 2, first author, 1 corresponding author]
    Short Answer:---------------------

    If your response to previous question was "No", which journals have you prominently published in? Please name no more than 5. Please avoid ambiguous abbreviations.

22. Did you have any preprints (which were not yet accepted/published at the time of application)? If yes, how many? (Suggested format: yes/no, number)
    Short Answer:---------------------

23. How many preprints have you posted throughout your career?
    Short Answer:---------------------

24. Do you have any patents filed (approved or pending)? If yes, how many? (Suggested format: yes/no, number)
    Short Answer:---------------------

25. Do you have any teaching experience?
    No
    Yes TA position(teaching assistantship)
    Yes experience beyond TA

26. If "yes, beyond TA", please elaborate: (e.g. What type of experience? How many years? Was it an undergraduate or graduate course? Was the course designed by you?)
Long Answer:---------------------

27. Did you have a PhD or postdoc fellowship?
No
Yes, PhD fellowship
Yes, postdoc fellowship
Yes, both
Yes

28. Have you ever been a PI or co-PI on a grant (such as K99/K01/R01 in the US, i.e. not postdoc/training fellowships)?
No
Yes, PI
Yes, co-PI
Other: Short Answer---------------

29. If you said "Other" to previous question or would like to name the type of grant please explain.
Long Answer:---------------------

30. Did you also apply for non-faculty positions such as Industry or Government or other jobs)?
No
Yes, non-faculty positions in academia
Yes, positions outside of academia

31. If you said "Other" to the previous question or would like to explain your answer please elaborate. Long Answer:---------------------

32. Do you have any comments? For example, was any aspect of your career particularly helpful/an obstacle when applying (preprints, grants etc...)?
Long Answer:---------------------

#### The Search committee survey

**Survey of faculty members involved in tenure-track searches**

Please answer in as much detail as you can. All responses will be treated fully anonymously.
*Required

1. How would you broadly define the field of the search(es) you've been involved in? *
   Biological and/or biomedical/life sciences
   Engineering
   Computer Science
   Mathematics
   Physics
   Chemistry
   Psychology
   Social Sciences
   Other

2. In what country are you based at? Short Answer:---------------------

3. How would you describe your institution? *
   PUI (Primarily Undergraduate Institution)
   D/PU: Doctoral/Professional Universities
   R1: Doctoral Universities
   R2: Doctoral Universities
   Independent research institute
   Prefer not to disclose
   Other: Short Answer---------------

4. Approximately how many applicants for a posted position do you get? *
   1-19
   20-49
   50-99
   100-199
   200+
   Prefer not to disclose

5. Approximately how many applicants make it through the first round of cuts? *
   1-19
   20-39
   40-59
   60+
   Prefer not to disclose

6. Approximately how many applicants are invited for off-site interview (Skype/phone)? *
   Short Answer---------------

7. Approximately how many applicants are invited for on-site interview ? *
   Short Answer---------------

8. Approximately how many offers does your committee make per job posting? *
   0-1
   2-3
   4+
   Prefer not to disclose

9. Approximately how many openings has your department had in the last five years? *
   0-1
   2-3
   4-5
   6+

10. Does your committee weigh Cell, Science, or Nature papers above papers in other journals? *

    Not at all Heavily

    1 2 3 4 5

11. To what extent does journal impact factor explicitly weigh in to the selection process (e.g. does the word ‘impact factor’ come up in discussions around applicants)? *

    Not at all Heavily

    1 2 3 4 5

12. To what extent does good mentorship in the candidate's postdoctoral/graduate student lab explicitly weigh on selection process (e.g. "This candidate's mentor is known to produce good trainees")? *

    Not at all Heavily

    1 2 3 4 5

13. To what extent does the research proposal weigh on the selection process (e.g. "This candidate's research statement is incredibly compelling!")? *

Not at all Heavily

1 2 3 4 5

14. Does your committee look favorably upon preprints? *

Yes (preprints are appreciated and considered a demonstration of productivity)

No (preprints are largely ignored)

Other: Short Answer---------------

15. How heavily does the committee weigh graduate student fellowships or awards (e.g. NSF GRF, NIH F30, etc.)? *

Not at all Heavily

1 2 3 4 5

16. How heavily does the committee weigh non-transitional postdoctoral fellowships or awards (e.g. NIH F32, AHA etc.)? *

Not at all Heavily

1 2 3 4 5

17. How heavily does the committee weigh transition awards as a positive factor (i.e. K99/R00 award, Burroughs Wellcome Career Award, or some award that provides the applicant with money as a new faculty member)? *

Not at all Heavily

1 2 3 4 5

18. How heavily does the committee weigh prior teaching experience? *

Not at all Heavily

1 2 3 4 5

19. What is your perception of the job market for tenure track faculty as someone involved in the search process (please tick all that are true). *

    Easy to identify good candidates from applications
    Hard to identify good candidates from applications
    Too many good applicants
    Too few good applicants
    Applicants lack sufficient teaching experience
    Candidates surpass expectations during interviews
    Candidates fall below expectations during interviews
    The market has changed a lot since I applied for a faculty position
    Other: Long Answer---------------

20. What information do you wish more candidates knew when they submit their application?
    Long Answer:---------------------

21. How long have you been involved in academic search committees? *
    1 - 4 years
    5 - 10 years
    11 - 19 years
    20 - 29 year
    30+ years
    Prefer not to disclose

22. Have you noticed any changes in the search process since the first search you were involved in?
    Long Answer:---------------------

23. Do you have any other comments or thoughts about the state of hiring for tenure track positions?
    Long Answer:---------------------
